# GABA Neurons in the Amygdala play a central Role in Urocortin-3–Mediated Stress Suppression of Reproduction

**DOI:** 10.1101/2025.01.06.631361

**Authors:** Junru Yu, Saeed Farjami, Kateryna Nechyporenko, Xiao Feng Li, Hafsa Yaseen, Yanyan Lin, Jinbin Ye, Owen Hollings, Ross de Burgh, Baban Singh, Kevin T. O’Byrne, Krasimira Tsaneva-Atanasova, Margaritis Voliotis

## Abstract

Stress can disrupt menstrual cycles, cause infertility, and lead to other reproductive disorders. The posterodorsal medial amygdala (MePD) processes stress signals and regulates the gonadotropin-releasing hormone (GnRH) pulse generator through GABAergic inhibitory projections to the hypothalamus. However, how stress is processed in the MePD - especially involving its dense GABA and Urocortin-3 (UCN3) neurons - remains poorly understood.

In this study, we combine *in vivo* GRadient-INdex (GRIN) lens mini-endoscopic calcium imaging (to track neuronal activity), optogenetics, clustering analysis, and computational modeling to investigate the MePD circuitry. Our findings reveal two anti-correlated GABA subpopulations in the MePD that dictate responses to both UCN3 neuron stimulation and restraint stress. Our computational modeling suggests that mutual inhibition between these GABA groups drives the anti-correlated activity and predicts how these interactions shape downstream responses to stimulation of GABA and UCN3 neurons.

We test these predictions using optogenetics and confirm that GABA neurons are critical for the transmission of UCN3 signals to regulate luteinizing hormone (LH) pulse frequency. Our study is the first to show how GABA neurons in the amygdala mediate stress effects on reproductive health, uncovering key neural mechanisms linking emotional and reproductive functions.

## Introduction

Successful reproduction requires normal function of a neural oscillator, the gonadotropin-releasing hormone (GnRH) pulse generator, that drives the pulsatile release of gonadotropic hormones, luteinizing hormone (LH) and follicle stimulating hormone (FSH) to control ovulation and spermatogenesis. This oscillator comprises Kisspeptin neurons co-expressing Neurokinin-B (NKB) and Dynorphin A (acronym KNDy) in the hypothalamic arcuate nucleus (ARC). The KNDy neural network not only provides the essential episodic stimulatory kisspeptin signal to the GnRH neurons, but receives diverse afferent inputs and hence is a hub for integrating various internal and external cues that regulate reproductive function^1,2^. Despite the marked heterogeneity of the myriad stressors, including acute versus chronic experimental stress paradigms, the common denominator is suppression of GnRH pulse generator frequency, which is associated with stress-related reproductive disorders and infertility^3–8^.

The amygdala, a part of the limbic brain typically associated with emotions and anxiety, has strong projections to the KNDy system^1,2^. The medial amygdala, and more specifically its posterodorsal subnucleus (MePD), has been shown to be a robust upstream regulator of GnRH pulse generator frequency^9–11^, pubertal timing^12,13^, sociosexual behavior^14,15^, anxiety^14^ and psychological stress-induced suppression of pulsatile LH secretion^16,17^.

In the MePD, the majority of neurons are GABAergic, accounting for a substantial portion of its projections, many of which target important hypothalamic centers involved in reproduction^18–20^, including the KNDy neurons and optogenetic stimulation of the latter suppresses LH pulsatility^17^. Moreover, chemogenetic inhibition of GABA neurons in the MePD can reverse the inhibitory effect of psychogenic stress on LH pulsatility^17^, suggesting an involvement of GABA signaling in the stress-induced suppression of pulsatile LH secretion. Although the precise way in which the MePD GABA neurons influence LH pulsatility is not fully understood, it has been suggested that they interact with the stress neuropeptide urocortin-3 (UCN3) neurons to modulate or reinforce stress-induced suppression of LH pulse frequency^16^.

UCN3, part of the corticotropin-releasing factor (CRF) superfamily and selective for CRF type 2 receptors (CRFR2)^21^, is highly expressed in the MePD^16,22^. Furthermore, UCN3 neurons in the MePD mediate psychosocial stress-induced suppression of the GnRH pulse generator and corticosterone secretion, thus playing a central role in the interaction between the reproductive and stress axes^16,23^. Previously we have shown that intra-MePD GABA antagonism can reverse UCN3-induced LH pulse suppression^23^, suggesting that GABA acts downstream of UCN3 signaling within the MePD. However, the local neural circuitry within the MePD mediating stress related information to the GnRH pulse generator remains to be established.

Here, we employed GRadient INdex (GRIN) lens mini-endoscopy to monitor and compare the calcium activity of GABA neurons at single-cell resolution under acute psychological stress or optogenetic stimulation of UCN3 neurons. Clustering analysis of the observed calcium transients uncovered two distinct GABA subpopulations exhibiting anti-correlated activity. To explore the functional role of these two subpopulations and unpick the mechanisms driving their anti-correlated activity, we build upon our previously developed MePD circuitry mathematical modeling framework^11^. Our modeling reveals that mutual inhibitory interactions between the GABA populations are sufficient to drive anti-correlated activity and predicts a complementary inhibitory effect of GABA and UCN3 signaling on the frequency of the GnRH pulse generator, which we tested by applying Cre/Flp recombinase intersectional strategies to selectively target and manipulate heterogeneous MePD neuronal populations^17^. Hence, by uncovering precise interactions between distinct subpopulations of MePD GABA neurons and UCN3 neurons in transmitting stress signals to the GnRH pulse generator, we provide critical and previously unknown insights into the neural circuitry driving stress-induced reproductive dysfunction. This discovery significantly advances our understanding of how emotional stress directly impacts reproductive health and identifies potential neural targets for therapeutic intervention in stress-related reproductive disorders.

## Results

### Dynamic reconfiguration of MePD GABA neuronal networks during selective optogenetic stimulation of urocortin-3 (UCN3) neurons and restraint stress

To investigate how the posterodorsal subnucleus of the medial amygdala (MePD) processes stress signals and to understand the role of GABA and UCN3 neurons in this process, we employed *in vivo* calcium imaging using GRadient INdex (GRIN) mini-endoscopy. A cohort of six female UCN3Cre::VGAT-Flpo transgenic mice were injected with a viral mixture containing pAAV-Syn-Flexrc[Chrimson-tdTomato] and pAAV-EF1a-fDIO-GCaMP6s. This intersectional viral strategy enabled dual functionality: (i) optogenetically stimulating MePD UCN3 neurons via Chrimson; and (ii) detecting calcium activity in MePD GABA neurons using GCaMP6s (Fig. 1**A**). GRIN lenses were surgically implanted at the same location as the viral injection, enabling imaging of calcium activity with single-neuron resolution Fig. 1**B-G**). The specificity of the AAV-Syn-Flex-rc[Chrimson-tdTomato] expression in the MePD was quantified by overlapping analysis with the UCN3 immunohistochemical staining and revealed that 95.06 ± 2.24% of tdTomato-positive neurons co-localized with the UCN3 immunohistochemical positive neurons (Supplementary Fig. 1). Further, 81.61 ± 6.03% of the total UCN3-immunoreactive neurons co-expressed Chrimson-tdTomato (Supplementary Fig. 1).

**Figure 1:**
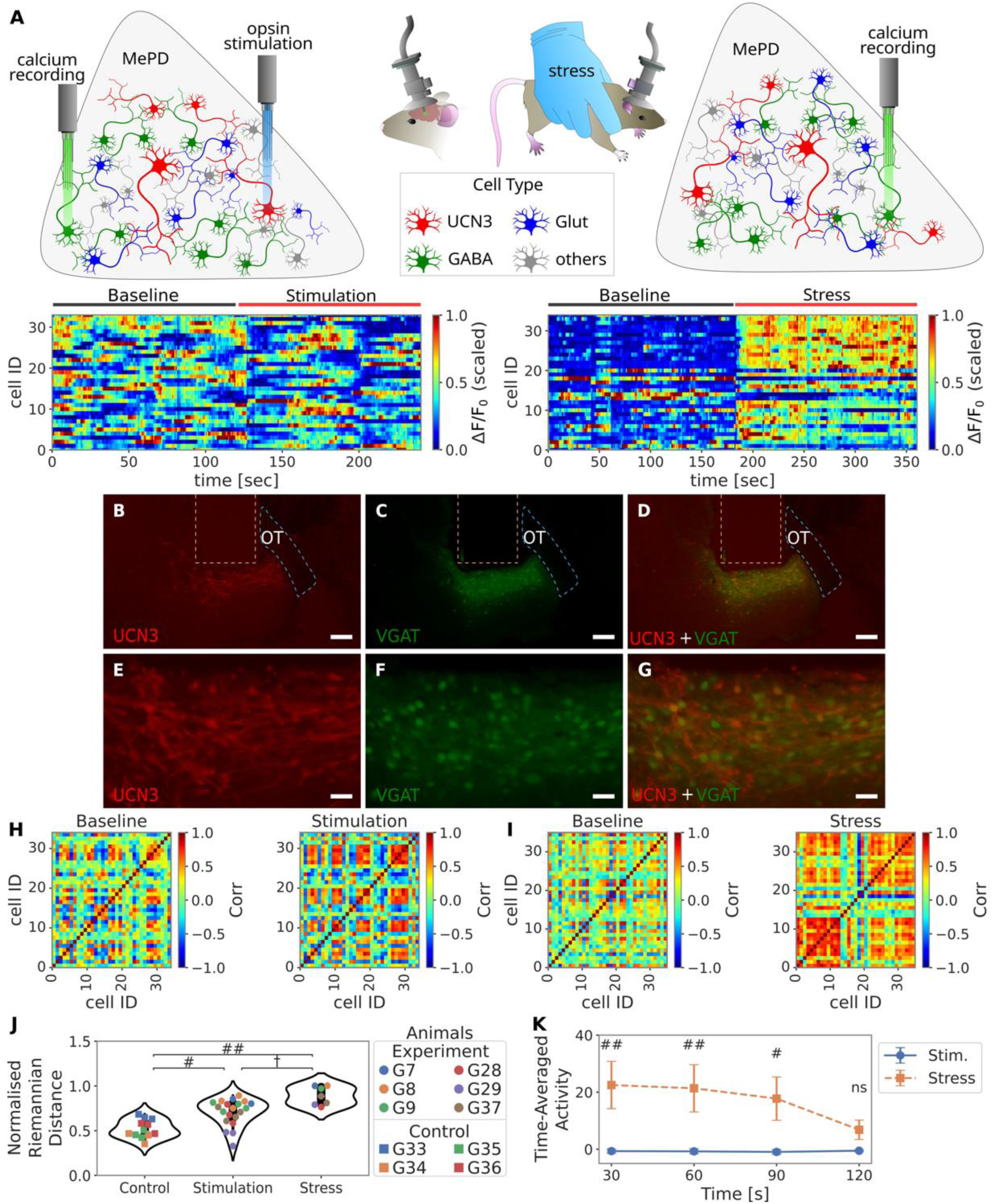
Functional connectivity of GABA neurons in posterodorsal medial amygdala (MePD) reconfigures in response to optogenetic stimulation of urocortin-3 (UCN3) neurons or acute restraint stress. (**A**) Schematic diagram of the experimental design for stimulating MePD UCN3 neurons and simultaneously imaging calcium activity from MePD GABA neurons. Representative heatmaps of calcium activity (ΔF/F_0_) recorded from MePD GABA neurons (left) in response to optogenetic stimulation of UCN3 neurons, (animal G7 trial 2, 34 neurons, see Supplementary Table 1; GABA neurons meeting our criteria for UCN3 expression excluded see Supplementary Fig. 2) and (right) during restraint-induced stress (animal G28, 35 neurons, see Supplementary Table 2). **(B-G)** Representative photomicrographs of the MePD from a UCN3-Cre::VGAT-Flpo mouse injected with pAAV-Syn-Flex-rc+pAAV-EF1a-fDIO-GCaMP6s. (**B**&**E**) Red fluorescence (tdTomoato) labels UCN3 neurons expressing Chrimson-tdTomato, (**C**&**F**) green fluorescence labels GABA neurons, and (**D**&**G**) red and green signals merged. (**H**&**I**) Functional connectivity matrices during baseline and under UCN3 activation and stress were constructed by calculating the pairwise Pearson’s correlation between calcium time-traces. (**J**) Normalized Riemannian distance between the correlation matrices from baseline and stimulation/stress periods. Each data point represents a single experiment, with different animals color-coded; control animals are shown as squares, and channelrhodopsin-expressing animals as circles. (**K**) Difference in time-averaged calcium activity between periods of UCN3 stimulation or stress and the baseline period. Scale bars: **B**-**D**, 200 µm; **E**-**G**, 50 µm.

Two series of experiments were conducted to assess GABAergic responses under distinct conditions. First, to assess the role of UCN3 neurons in recruiting GABAergic response, baseline calcium activity from MePD GABA neurons was recorded for 3 min, followed by 3 min of recording during optogenetic stimulation of UCN3 neurons (Fig. 1**A**). Optogenetic stimulation was delivered in a discontinuous pattern, 5 seconds on, 5 seconds off, with a light-pulse frequency of 10 Hz during the ‘on’ periods (10ms pulse width, 10 mW power, and 532 nm wavelength). GABA neurons with activity that mimics the pattern of optic stimulation, evidence that they are co-expressing UCN3, are removed from further analysis (see Methods and Supplementary Fig. 2). To further characterize the GABAergic response to acute stressors, baseline calcium activity of GABA neurons was recorded for 2 min, followed by 2 min of recording while the animals were exposed to restraint stress (Fig. 1**A**). The dynamic GABAergic response to optogenetic stimulation of UCN3 neurons and restraint stress was summarized in functional terms using pairwise (neuron-neuron) association matrices, constructed from Pearson correlation coefficients between neuronal trace pairs (see Methods). These correlation matrices represent the MePD GABAergic response as a graph of interconnected nodes (neurons) with edge weights corresponding to the strength of their pairwise functional interactions (Fig. 1**H**&I)^24^. To quantify the effect of stress or UCN3 activation on functional connectivity we calculated the normalized Riemannian distance between the correlation matrices from baseline and stimulation/stress periods. Our analysis revealed significant alterations in the functional connectivity of the GABAergic network during both UCN3 stimulation and stress relative to control animals lacking channelrhodopsin expression (Fig. 1**J**). Furthermore, on average, stress induces a significantly higher population time-averaged calcium response than UCN3 stimulation; however, this effect diminishes over the 2-minute stress period (Fig. 1**K**). These results indicate that the GABA population in the MePD plays a pivotal role in processing stress signals, such as those induced by restraint stress. Furthermore, our findings suggest that UCN3 neurons may serve as a key relay, driving functional reorganization within the GABAergic population to support transmission of stress-related information.

### Clustering analysis reveals two functionally distinct GABA subpopulations in the MePD

To better understand the functional organization of the GABA MePD network during optogenetic stimulation of UCN3 neurons and under restraint stress, we performed clustering analysis on the functional connectivity matrices we obtained from the calcium recordings of GABA neurons.

Clustering results consistently revealed two functionally distinct cell populations both during optogenetic stimulation of UCN3 neurons and restraint stress (Fig. 2**A&C**). Under both conditions, activity between the two clusters appeared highly anti-correlated as illustrated by two representative examples depicted in Fig. 2**B&D**. Moreover, we find that clusters are significantly different in size, with the smallest cluster accounting on average for 35% of all cells (Fig 2**E&F**, Supplementary Fig. 3). The robustness of the above finding was also verified using the K-means clustering algorithm and an alternative measure of association between neuronal traces that accounts for time-lag and sign in neuronal interactions (see Methods and Supplementary Figs. 4-6).

**Figure 2:**
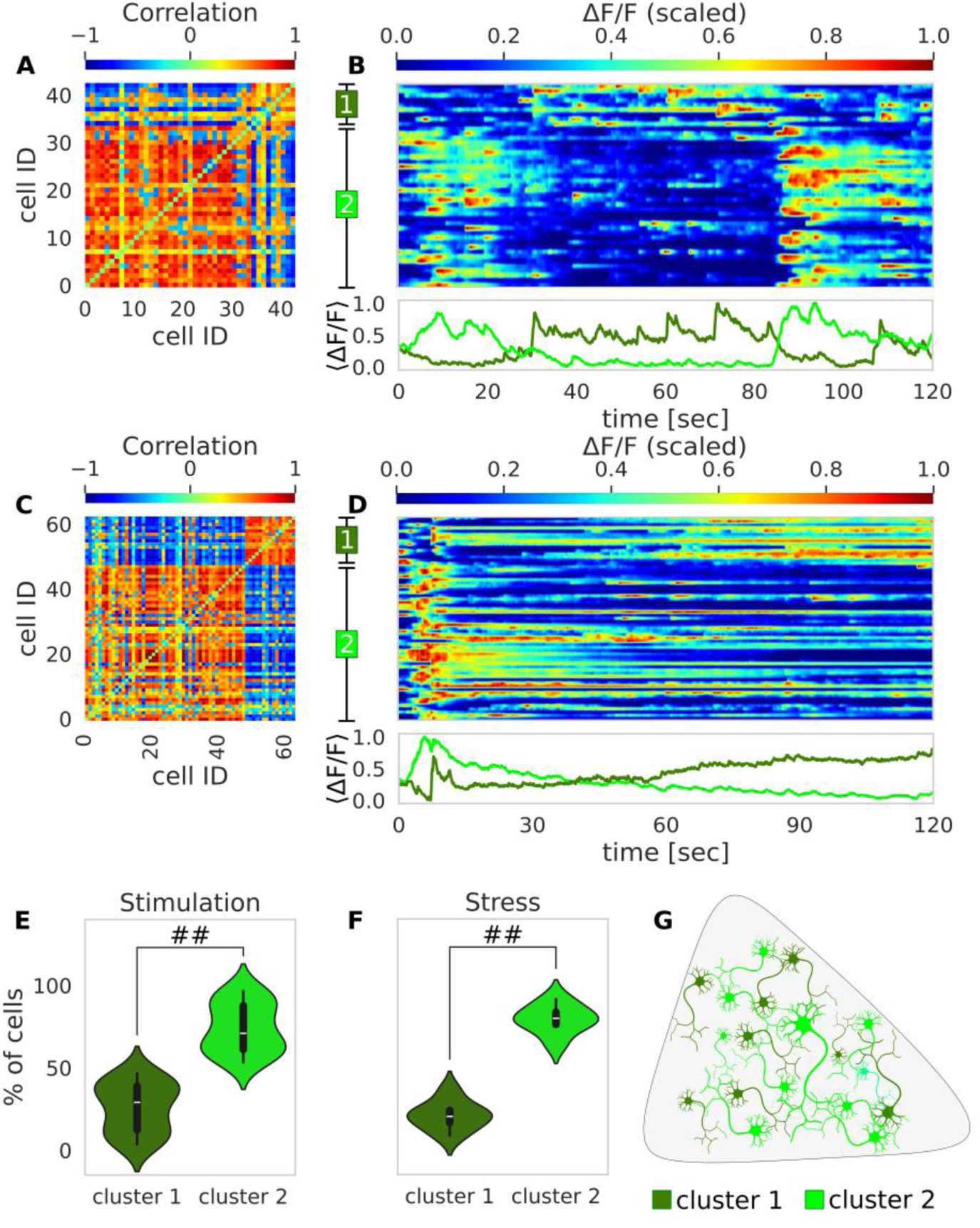
Clustering identifies two functionally distinct GABA subpopulations in the posterodorsal medial amygdala (MePD) both during UCN3 optogenetic stimulation and restraint stress. Representative example of (**A**) a clustered connectivity matrix and (**B**) associated calcium time-traces of MePD GABA neurons during optogenetic stimulation of UNC3 neurons and the population average below. Representative examples of (**C**) a clustered connectivity matrix and (**D**) associated calcium activity time-traces of MePD GABA neurons during restraint stress and the average population activity below. (**E**&**G)** Relative sizes of the two GABA subpopulations identified via clustering analysis. (##: p<0.001).

Varying the frequency of optogenetic stimulation of UCN3 neurons influenced the functional connectivity within the GABA network (Supplementary Fig. 7). Stimulation at 5 Hz produced no significant change compared with controls, 10 Hz resulted in a statistically significant increase, whil 20 Hz reduced the effect, which could possibly reflect a differential release of neurotransmitters/neuropeptides at higher frequencies^25^. In contrast, stimulation at 5, 10, or 20 Hz did not significantly affect the correlation between the average activity of the two GABA clusters (Supplementary Fig. 8). Based on the functional role of the MePD in relaying stress signals to the hypothalamic arcuate nucleus (ARC), our results suggest that the identified GABA subpopulations (clusters) could correspond to distinct populations of GABA interneurons and GABA efferent neurons—defined as neurons that transmit signals from the MePD to downstream targets. Known efferent projections to the ARC play a role in regulating the gonadotropin-releasing hormone (GnRH) pulse generator, a key driver of reproductive function^11,17^.

### Mutually inhibitory interactions between GABA populations drive anti-correlated activity

To explore the network mechanisms driving the anti-correlated activity of the GABA subpopulations, we build upon our previously developed MePD circuit model^11^. The model allows us to explore interactions between MePD neuronal populations and understand emergent patterns of dynamical activity. Specifically, the model describes the calcium dynamics of two interacting populations of GABA neurons, corresponding to interneurons and efferent projection neurons. Efferent neurons project their axons outside the MePD to downstream targets such as the KNDy network. Both GABA populations interact with a glutamatergic population (Fig. 3**A**). Experimental evidence shows that antagonizing glutamate signaling completely blocks the suppressive effect of MePD UCN3 activation on LH pulsatility^23^, highlighting a functional role of glutamatergic neurons in the circuit. Therefore, the activity of the glutamatergic population is also included in the MePD model alongside the two GABA populations. Model equations are provided in Supplementary Information, and the parameter values for the MePD model are in Supplementary Table 3.

**Figure 3:**
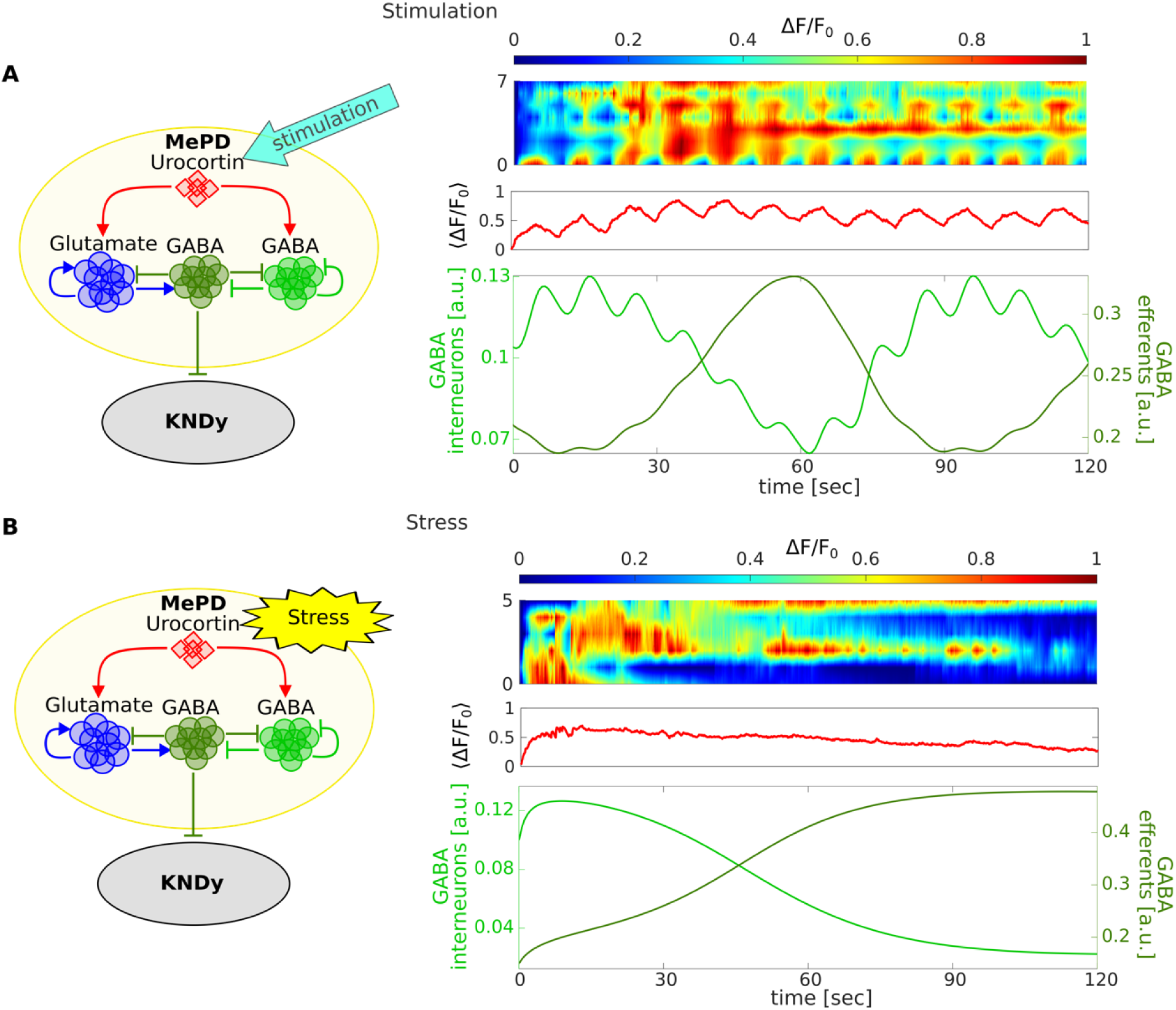
Mathematical model simulating the activity in the GABA neuronal populations in the MePD under the increased excitatory input from UCN3 neurons and mimicking the conditions under restraint stress. The schematics on the left provide a visual representation of the assumptions built into our mathematical model of the MePD neuronal circuit, consisting of two GABA populations and a glutamatergic population, that interact with UCN3. **(A)** Calcium time-traces of MePD UCN3 neurons during optogenetic stimulation of UCN3 neurons and corresponding mean activity. Model simulations of the effects of UCN3 optogenetic stimulation supports the emergence of anti-correlated activity in GABA subpopulations. **(B)** Calcium time-traces of MePD UCN3 neurons during restraint stress and corresponding mean activity of the UCN3 calcium traces during restraint stress with corresponding mean activity. Simulations of the MePD circuit under the effects of stress also support anti-correlated GABA subpopulations’ activity.

To incorporate input from UCN3 neurons we include a non-autonomous term in the model informed by recordings of the activity of UCN3 neurons during optogenetic stimulation (see Fig. 3 **A** and Methods). The recorded traces exhibit a periodic pattern driven by the experimental stimulation protocol with 5 seconds on, 5 seconds off (see Methods). In the model, we approximated this UCN3 input using a constant drive along with a sinusoidal function with roughly 5 seconds on, 5 seconds off pattern, driving the populations GABA interneurons and the glutamatergic neurons. To simulate the anti-correlated activity between the two GABA neuronal subpopulations while also reproducing previously published results on the effects of neuropharmacological perturbations in the MePD on LH pulsatility^23^, we find that we need to include bi-directional (i.e. mutually inhibitory) GABA interaction in the circuit as well as GABA interneurons self-inhibition (Fig. 3**A**).

Next, we validate our model by simulating the effects of stress on the MePD circuit. UCN3 neurons in the MePD have also been shown to mediate psychosocial stress-induced suppression of the GnRH pulse generator^16^. Experimentally recorded calcium activity of UCN3 neurons during stress intervention is characterized by a fast rise followed by a slow decay (Fig. 3**B**), which we model by taking the difference between two exponential functions, with one fast and one slow time constant (see Methods). Using this input, our mathematical model confirms the emergence of anti-correlated behavior between the GABA subpopulation whereby the activity of GABA interneurons decreases over the stress period, while the activity of the GABA efferent neuron population increases (Fig. 3**B**), supporting the proposed MePD circuit structure (Fig. 3**A**).

### *In silico* modeling reveals complementary relationship between MePD GABA and UCN3 populations

Having shown that the model of the MePD circuitry reproduces the observed anti-correlated dynamics, we next couple it to our network model of the KNDy population^11,26^. KNDy model details are in Supplementary Information, and the model parameter values can be found in Supplementary Table 4. Since KNDy neuronal activity serves as a well-established proxy for LH secretion, we use the dynamics of the KNDy network model to infer changes in LH pulsatility in response to MePD circuit manipulations. Our previous in silico findings^11^ show that MePD signal integration is predominately mediated by GABA projections. The effects of MePD GABA and glutamate antagonism alone and together with UCN3 stimulation on LH pulsatility have been previously investigated^23^, and we use these experimental findings to calibrate our coupled model parameters by reproducing these effects accordingly (see Supplementary Fig. 9).

Next, using the calibrated model, we predict that providing a periodic excitatory input (see Supplementary Fig. 10**A**), that mimics the effects of optogenetic stimulation of UCN3, leads to the increase in the KNDy inter-pulse interval (IPI) from 15.18 min to 23.97 min due to increased activity in the inhibitory GABAergic input to the KNDy network (Fig. 4**A**).

**Figure 4:**
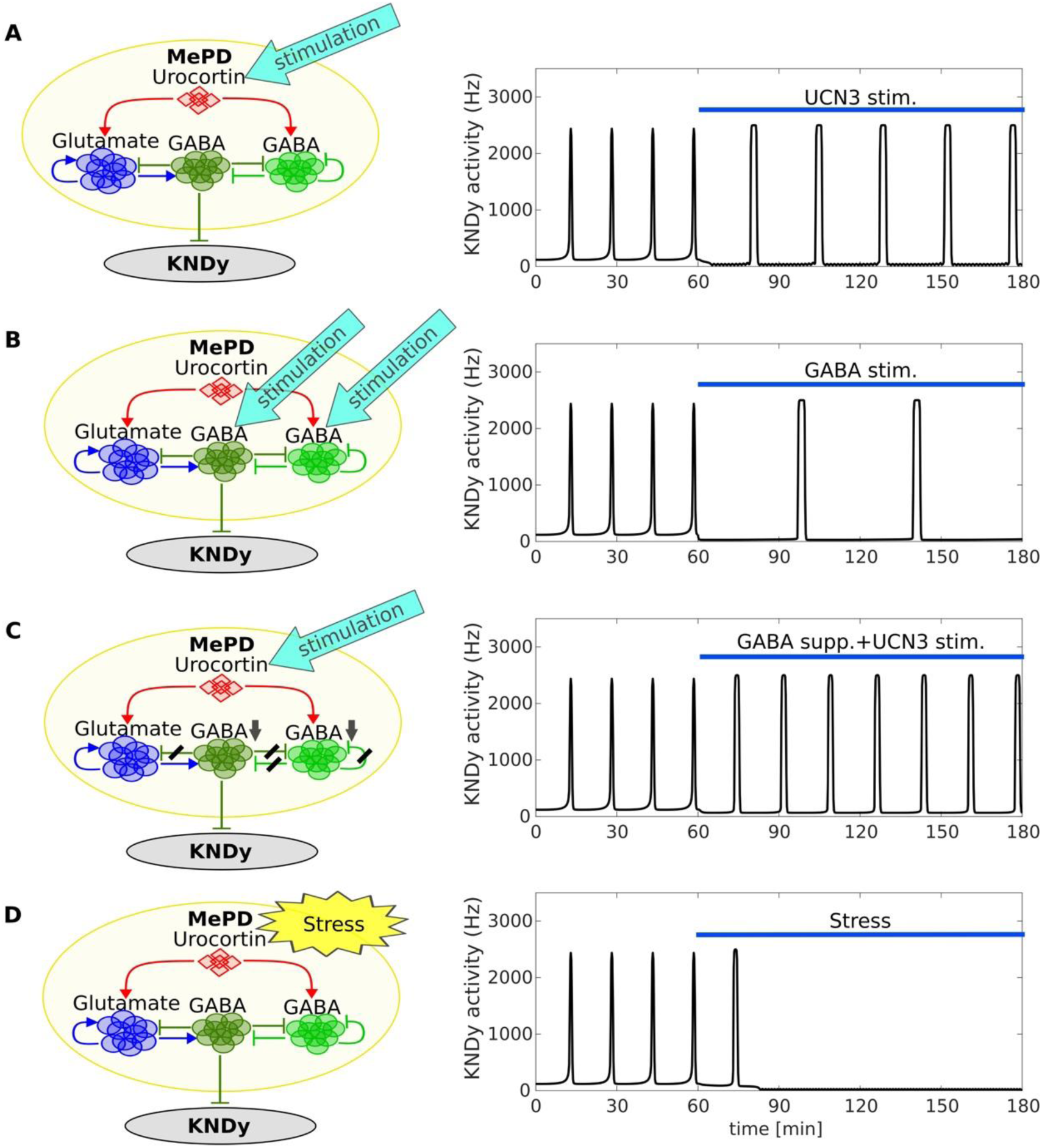
Coupled MePD - KNDy model predictions for the effect of stress and stimulation of UCN3 neurons on GnRH pulse generator activity and hence LH pulsatility. (**A**) Simulation of the optogenetic stimulation of the UCN3 neurons leads to the increase of the KNDy IPI 15.18 min to 23.97 min. (**B**) Mimicking the effects of the optogenetic stimulation of the MePD GABA neurons *in silico* (providing excitatory input to both GABA populations) increases KNDy IPI from 15.18 min to 42.65 min. (**C**) Simulation of the effects of the optogenetic stimulation of the UCN3 neurons and simultaneous inhibition of GABA neuronal populations do not significantly affect KNDy IPI, where it changes from 15.18 min to 17.32 min. (**D**) Simulating the effects of stress leads to the cessation of the pulsatile dynamics.

Since MePD has an overwhelming number of GABAergic outputs and interneurons, we are interested in their role in information processing of stress signaling. To this end we provide periodic excitatory input directly to both populations of GABA neurons (see Methods and Supplementary Information). This is different to the indirect activation through UCN3 stimulation, as it leads to the increased activity in both GABA subpopulations (Supplementary Fig. 10**B**). The resulting effect on the KNDy IPI is an increase from 15.18 min to 42.65 min (Fig. 4**B**).

In the model we hypothesize that UCN3 signaling to the KNDy is operating through the GABAergic pathway, hence disrupting GABAergic interactions will suppress UCN3’s effects on KNDy. To investigate it we suppress the strength of the GABAergic interactions by 50% and mimic UCN3 stimulation by providing excitatory input to the circuit. The model shows that this intervention does not have a significant effect on KNDy IPI, apart from slightly increasing it from 15.18 min to 17.31 min (Fig. 4**C** & Supplementary Fig. 10**C**).

Taken together the above could be summarized as the following set of model predictions along with suggested ways to test them experimentally:

1. Optogenetic stimulation modelled via excitation of the MePD UCN3 increases the KNDy IPI 15.18 min to 23.97 min. This could be tested experimentally via in vivo optogenetic activation of the MePD UCN3 neuronal population and simultaneous measurement of LH pulsatility.
2. Stimulation of the MePD GABA populations increases the KNDy IPI 15.18 min to 42.65 min. Such a substantial change underscores the critical role of GABAergic signaling in modulating KNDy pulsatility. Experimental confirmation could involve in vivo optogenetic activation of the entire MePD GABA population accompanied by LH pulsatility measurement.
3. Stimulation of the MePD UCN3 with simultaneous suppression of the GABAergic interactions in the MePD does not significantly affect KNDy IPI, changing it from 15.18 min to 17.31 min. This finding suggests that UCN3 effects on the KNDy network are primarily dependent on its interactions with GABA populations. A strategy to verify this hypothesis experimentally could be to stimulate UCN3 non-GABA-expressing neurons in vivo while simultaneously inhibiting non-UCN3 GABA neurons and measure LH pulsatility.

Finally, we investigate in silico the effects of stress on LH pulse dynamics mediated via the MePD. In the restraint-stress experiment it was shown that one GABA cluster was active, while the other one was inactive (Fig. 2**D**). By simulating these dynamics (Supplementary Fig. 9**D**) in the coupled MePD-KNDy network model we are able to demonstrate that pulses in the KNDy network ceased (Fig. 4**D**). These modeling findings align with published experimental evidence indicating that restraint stress suppresses LH pulsatility^16^.

### Selective optogenetic stimulation of UCN3 non-GABA neurons inhibits pulsatile LH secretion in female mice

We now test experimentally our modeling predictions detailed in the previous section regarding the inhibitory effect of MePD UCN3 activation on LH pulsatility in vivo in freely behaving mice. To this end a cohort of UCN3-Cre::VGAT-Flpo transgenic female mice was injected with an AAV-nEF-Con/Foff 2.0-ChRmine-oScarlet viral construct. This intersectional approach enabled conditional expression of channelrhodopsin (ChRmine) in UCN3 neurons not expressing GABA (UCN3 non-GABA) (Con/Foff group). Following the viral injection, a unilateral fiber-optic cannula was implanted into the MePD, above the injection site, to facilitate selective activation of the targeted neuronal population. The specificity of the AAV-nEF-Con/Foff 2.0-ChRmine-oScarlet expression in the MePD was quantified by overlapping analysis with the UCN3 immunohistochemical staining and revealed that 98.92 ± 1.08% of the oScarlet-positive neurons co-localized with the UCN3 immunohistochemical positive neurons (Supplementary Fig. 11). Further, 54.18 ± 5.04% of the total UCN3-positive neurons coexpressed ChRmine-oScarlet (Supplementary Fig. 11).

The stimulation protocol consisted of a 60-min control period, followed by 60 min of optogenetic stimulation. Stimulation was delivered, as above, in a pulsed pattern (5 seconds on, 5 seconds off) with a pulse frequency of 10 Hz during the on period (10-ms pulse width, 10 mW power, 532 nm wavelength). Blood samples (5 µL) were collected every 5 min throughout the protocol and used to measure LH levels.

Statistical analysis indicated post-stimulation LH inter-pulse intervals differed across the two groups (F (2, 30) = 15.83, P < 0.0001; two-way ANOVA). In the Con/Foff group, the LH inter-pulse interval significantly increased after optogenetic stimulation compared to the control period (15.47 ± 0.91 min vs. 23.75 ± 1.07 min, mean ± SEM, n = 6, p<0.01; Fig. **5A&D**), validating the first of our modeling predictions. In contrast, the no-stimulation group, which received the Con/Foff viral construct without optogenetic stimulation, showed no significant changes in LH inter-pulse interval (15.84 ± 0.37 min vs. 16.68 ± 0.86 min, n = 6, p > 0.05; Fig. 5**B&D**). Similarly, in a separate cohort of mice injected with a control virus (DIO-EYFP), optogenetic stimulation resulted in no significant differences in LH inter-pulse intervals before and after stimulation (19.83 ± 1.02 min vs. 17.78 ± 0.93 min, n = 6, p > 0.05; Fig. **5C&D**). The LH pulse amplitude and mean LH levels were quantified, and data are provided in Supplementary Table 5.

**Figure 5:**
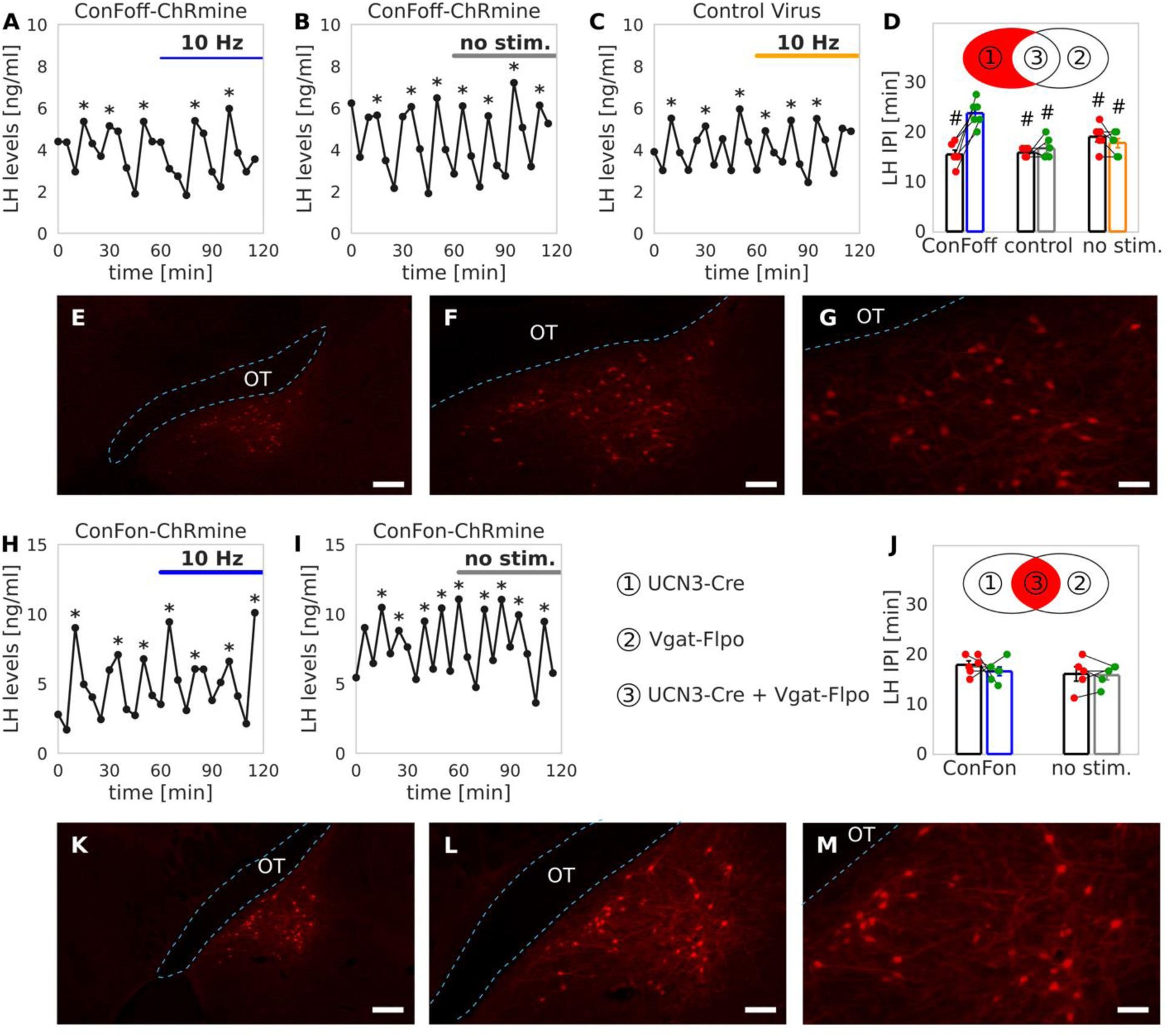
Effect of optogenetic stimulation of UCN3 non-GABA and UCN3 neurons co-expressing GABA in the MePD on LH pulse frequency. (**A**-**D**) Optogenetic stimulation of UCN3 non-GABA neurons inhibited LH pulses. Representative examples of LH pulsatility in UCN3-Cre::VGAT-Flpo mice injected with AAV-Con/FoffChRmine in response to (**A**) 10Hz optic stimulation or (**B**) no stimulation as control, and (**C**) in mice injected with control virus in response to 10 Hz stimulation. Pulses detected by the DynPeak algorithm are indicated with an asterisk (*). (**D**) Mean LH inter-pulse interval (IPI) (±SEM) for each group during the control period and over the subsequent stimulation period. #: p<0.01 vs stimulation period in Con/Foff group (two-way ANOVA, Tukey’s post-hoc). (**E**-**G**) Representative photomicrographs of expression of AAV-nEF-Con/Foff 2.0-ChRmine-oScarlet (red) in MePD UCN3 neurons. (**H**-**M**) Optogenetic stimulation of neurons co-expressing UCN3 and GABA in the MePD did not impact LH pulse significantly. Representative examples of LH pulsatility in UCN3-Cre::VGATFlpo mice injected with Con/Fon-ChRmine-oScarlet virus in response to (**H**) 10 Hz optic stimulation or (**I**) no stimulation as control. (J) Mean LH IPI (±SEM) for each group during the control period and over the subsequent stimulation period (two-way ANOVA). (**K**-**M**) Representative photomicrographs of neurons co-expression UCN3 and GABA, tagged with oScarlet (red). Scale bars represent (**E**&**K**) 200 µm, (**F**&**L**) 100 µm, (**G**&**M**) 50 µm. OT, optic track. The blue vertical bars in E to G and K to M indicate the position of the fiber optic cannula.

### Selective optogenetic stimulation of UCN3-expressing GABA neurons fails to affect pulsatile LH secretion in female mice

To further test the functional interactions between the UCN3 and GABA populations, a separate cohort of UCN3-Cre::VGAT-Flpo transgenic female mice was injected with an AAV-nEF-Con/Fon 2.0ChRmine-oScarlet viral construct, which allows optogenetic stimulation of UCN3 neurons coexpressing GABA (UCN3-expressing GABA) (Con/Fon group). The specificity of the AAV-nEFCon/Fon 2.0-ChRmine-oScarlet expression in the MePD was quantified by overlapping analysis with the UCN3 immunohistochemical staining and revealed that 98.77 ± 1.24% of the oScarlet-positive neurons co-localized with the UCN3 immunohistochemical positive neurons co-expressing GABA (Supplementary Fig. 12). Further, 32.73± 2.60% of the total UCN3-positive neurons co-expressed ChRmine-oScarlet (Supplementary Fig. 12).

Statistical analysis showed no significant interaction effect of group and optogenetic stimulation on LH inter-pulse intervals (F (1, 18) = 0.2709, p = 0.6091; two-way ANOVA). That is, LH inter-pulse intervals in the Con/Fon group, showed no change following optogenetic stimulation compared to the control period (17.92 ± 0.80 min vs. 16.60 ± 0.88 min, mean ± SEM, n = 6, p > 0.05; Fig. **5H&J**). Similarly, in the no-stimulation control experiment for this cohort, which did not receive optogenetic stimulation, there was no significant difference in LH inter-pulse intervals (16.08 ± 1.45 min vs. 15.83 ± 0.95 min, n = 5, p > 0.05; Fig. 5**I&J**). The LH pulse amplitude and mean LH levels were quantified, and data are provided in Supplementary Table 5.

Taken together, our results suggest that although UCN3 and GABAergic neuronal populations overlap, they serve distinct functional roles within the MePD circuit, with both being critical for relaying inhibitory signals to the hypothalamic GnRH pulse generator. While independent activation of either population inhibits the GnRH pulse generator, activation of UCN3 neurons that co-express GABA is insufficient to elicit this effect.

### The MePD GABA neurons relay information from UCN3 neurons and inhibit the GnRH pulse generator

To test our remaining model predictions and establish the functional role of GABA neurons in the MePD in transmitting inhibitory signals to the hypothalamic GnRH pulse generator, we injected the AAV-EF1a-DIO-ChR2-EYFP viral construct into VGAT-Cre transgenic mice, targeting the entire MePD GABA neuron population for expressing ChR2 and used the optogenetic stimulation protocol described above (60 min control period followed by 60 min stimulation) to stimulate the entire MePD GABA population. In the VGAT-Cre-tdtomato mice, we observed that 93.27 ± 2.99% of the EYFP positive neurons were tdtomato-positive and 87.01± 5.21% of the tdtomato-positive neurons were EYFP-positive. This indicates a high degree of co-expression consistency between the viral and transgenic reporters. Statistical analysis showed a significant interaction effect of our genetic intervention and stimulation period on LH pulsatility (F (2, 24) = 10.55, p = 0.0005; two-way ANOVA). Stimulation of the GABA population markedly increased the LH inter-pulse interval compared to the pre-stimulation control period (18.5 ± 2.03 min vs 49.5 ± 8.46 min, mean ± SEM, n = 5, p<0.01; Tukey’s post-hoc; Fig. **6A&D**), validating modeling prediction 2. In contrast, mice that did not receive optogenetic stimulation showed no change in LH inter-pulse interval (21.25 ± 3.79 min vs 19.33 ± 1.52 min, n = 5, p > 0.05; Tukey’s post-hoc; Fig. **6B&D**). Similarly, control virus (AAV-EF1a-DIO-EYFP) injections did not result in any significant difference in LH inter-pulse interval before and during stimulation (16.50 ± 0.67 min vs 18.00 ± 0.62 min, n = 5, p > 0.05; Tukey’s post-hoc; Fig. **6C&D**). The LH pulse amplitude and mean LH levels were quantified, and data are provided in Supplementary Table 6.

**Figure 6:**
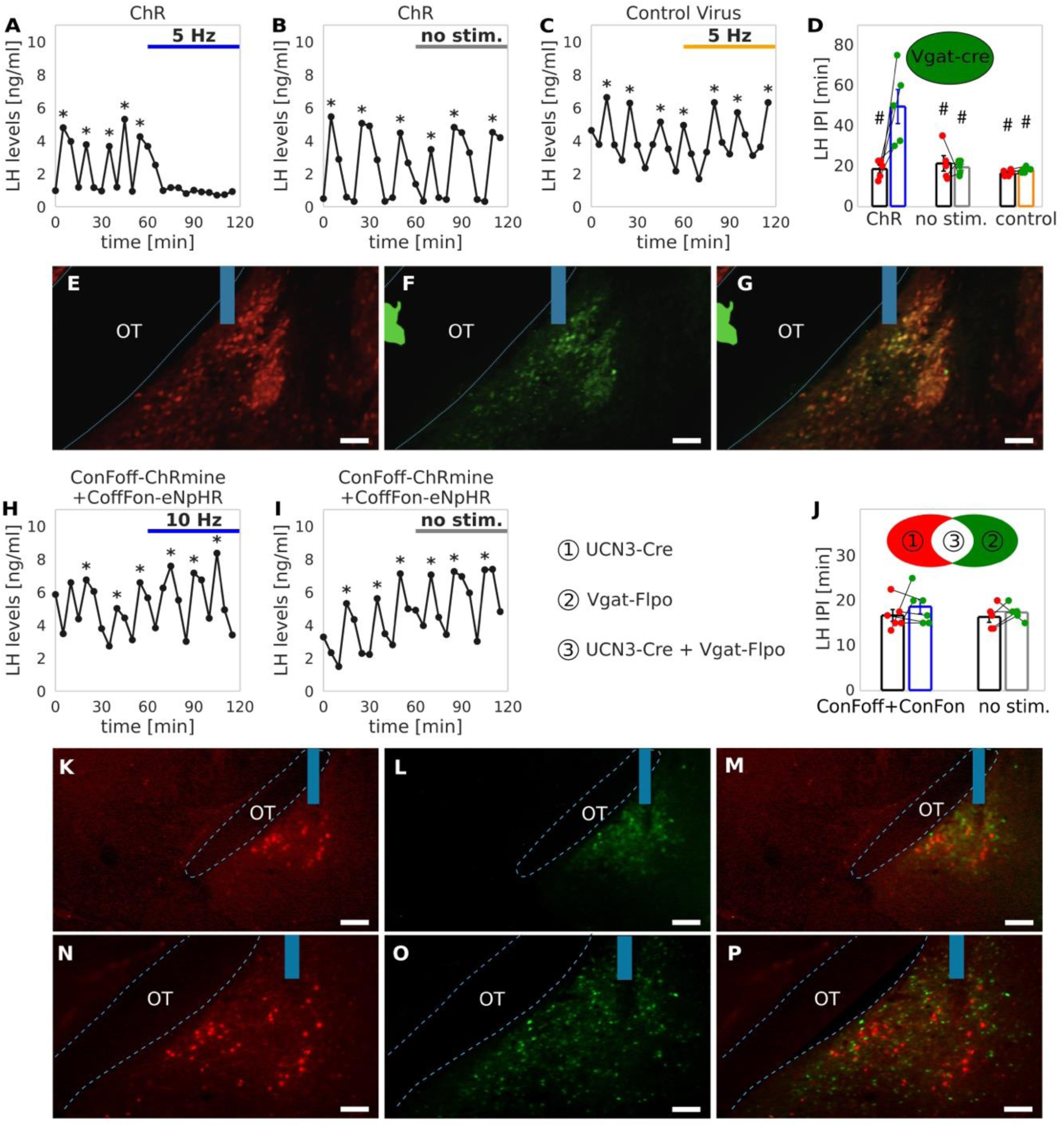
Selective optogenetic stimulation of GABA neurons in the MePD inhibits LH pulsatility and their activity is necessary for mediating UCN3 signaling downstream. (**A-G**) Optogenetic activation of the entire MePD GABA population in VGAT-Cre mice led to significant LH pulse suppression. Representative examples showing LH pulsatility in VGAT-Cre mice injected with AAV-double floxed-ChR2-EYFP virus in response to (**A**) 5 Hz stimulation or (**B**) no stimulation as control, and (**C**) 5 Hz stimulation in mice injected with control virus. (**D**) Mean LH IPI (±SEM) for each group during the control period and over the subsequent stimulation period. #: p<0.01 vs stimulation period in ChR2 group (two-way ANOVA, Tukey’s post-hoc). (**E-G**) Representative photomicrographs of MePD GABA neurons, labeled with (**E**) tdTomato (red), (**F**) ChR2-EYFP (green), and (**G**) merged. Scale bars represent. (**H-P**) The simultaneous inhibition of non-UCN3 GABA neurons in the MePD restored the LH suppression induced by stimulation of non-GABA UCN3 neurons. Representative examples showing LH pulsatility in UCN3-Cre::VGAT-Flpo female mice injected with Con/FoffChRmine-oScarlet and Coff/Fon-NpHR-EYFP virus mixture in response to (**H**) 10 Hz optogenetic stimulation or (**I**) no stimulation as control. (**J**) Mean LH IPI (±SEM) for each group during the control period and over the subsequent stimulation period. (two-way ANOVA). (**K-P**) Representative photomicrographs of fluorescence in MePD neurons. (**K**&**N**) non-GABA UCN3 neurons tagged with oScarlet (red), (**L**&**O**) non-UCN GABA neurons tagged with EYFP (green) (**M**&**P**) merged. Scale bars represent (**E-G**) 150 µm, (**K-M**) 200 µm, (**N-P**) 100 µm. OT, optic track. The blue vertical bars in **E** to **G** and **K** to **P** indicate the position of the fiber optic cannula.

Next, UCN3-Cre::VGAT-Flpo transgenic females were injected with an intersectional viral mixture containing AAV-nEF-Con/Foff 2.0-ChRmine-oScarlet and pAAV-nEF-Coff/Fon-NpHR3.3-EYFP, enabling simultaneous excitation of UCN3 non-GABA neurons and inhibition of non-UCN3 GABA neurons. Statistical analysis showed no significant interaction effect of genetic manipulation and stimulation period on LH pulsatility (F (1,18) = 0.131, p = 0.7216; way ANOVA). Moreover, optogenetic stimulation resulted in no significant change in the LH inter-pulse interval relative to the control period (16.67 ± 1.31 min vs 18.61 ± 1.58 min, mean ± SEM, n = 6, p > 0.05; Fig. **6H&J**), which aligns with our modeling prediction 3. Similarly, in mice injected with the same viral mixture but without receiving optogenetic stimulation, no significant change in LH inter-pulse interval was detected (16.33 ± 1.19 min vs 17.33 ± 0.81 min, n = 5, p > 0.05; Fig. **6I&J**). The LH pulse amplitude and mean LH levels were quantified, and data are provided in Supplementary Table 5.

Collectively, these findings suggest that MePD GABA neurons are integral to regulating LH pulse frequency. The inhibitory effect on LH pulses observed upon UCN3 neuron activation is attenuated when GABA neuron activity is optogenetically suppressed, implying that GABA signaling acts downstream of UCN3 neurons, relaying the inhibitory signal to the hypothalamic GnRH pulse generator.

## Discussion

The posterodorsal subnucleus of the Medial Amygdala (MePD) is a critical regulator of reproductive function. Single-cell transcriptomics has revealed a mosaic of glutamatergic and GABAergic populations in this region, with GABA neurons co-expressing receptors for gonadal steroids as well as neuromodulators and cognate receptors involved in parental and sociosexual behavior (including kisspeptin, prolactin and oxytocin)^27^. Furthermore, calcium imaging of neuronal ensembles from this region has started to decipher how this brain region encodes information about social interactions and stressful stimuli^28^, with stress-related information being relayed to the reproductive axis via potentially direct projections to the hypothalamus^1,2^, and eliciting well-documented responses in pubertal timing^12,13^, LH secretion^16,17^ and the GnRH pulse generator^9–11^. Here, to elucidate the neuronal circuitry involved in MePD processing of stress signals, we used GRIN lens mini-endoscopic calcium imaging to study, for the first time, the dynamic activity of MePD GABA neurons under acute restraint stress as well as the activity of GABA neurons in response to UCN3 activation. Functional network analysis of the imaging data revealed that acute restraint stress induces a reconfiguration of the GABA population, highlighting their role in encoding and transmitting stress information. Optogenetic stimulation of UCN3 neurons produced a comparable network reconfiguration indicating that both populations are co-involved in stress processing. The overall magnitude of calcium activity (ΔF/F₀) differed between the two conditions, with stress generally inducing a stronger response than optogenetic stimulation of UCN3 neurons. This discrepancy may reflect the engagement of additional stress-related pathways or neuromodulatory factors that are not fully replicated by isolated UCN3 stimulation, such as stress-induced GABA transmission that has been well-documented in the amygdala^17,23^. Clustering analysis of the calcium transients identified two distinct GABA subpopulations with anti-correlated activity. This finding of two distinct populations is consistent with physiological and morphological studies of GABA neurons in the medial amygdala characterizing them as either projecting cells or local^18,20^.

Building upon our previously developed computational model of the MePD circuit, we explored the network connectivity to understand the possible mechanisms through which anti-correlated activity emerges. The model consists of a glutamatergic subpopulation and two GABA subpopulations: one comprising GABA interneurons and the other GABA efferent projection neurons. Our analysis identified two conditions necessary for the emergence of anti-correlated activity between these GABA subpopulations, consistent with our experimental observations. First, mutual inhibition between the GABA subpopulations is required, such that activation of one population suppresses the other. In our mechanistic model, mutual inhibition arises from the biophysical characteristics of GABAergic transmission, where the release of GABA inhibits target neurons, thereby producing inhibitory interactions between the subpopulations. Second, if both subpopulations receive a shared input (e.g., from the glutamatergic population or any other population of excitatory cells), their dynamics become entrained to that common stimulus, meaning that distinct input streams to each subpopulation are necessary for the observed anti-correlation. To satisfy this condition in our model, GABA interneurons receive inputs from UCN3 neurons, whereas the GABA efferent projection neurons receive inputs from the glutamatergic population. This circuitry implies that stress information is processed along two key pathways (one glutamatergic and one GABAergic) and is integrated by the GABA efferent projection neurons to the arcuate KNDy system.

We note that an alternative connectivity pattern, with the UCN3 population directly projecting to the GABA efferent population, could also fit our experimental observations, but would not reproduce previously published results on the effects of neuropharmacological perturbations in the MePD on LH pulsatility^23^. Specifically, the suppression of glutamatergic neurons in combination with stimulation of UCN3 neurons. Under this scenario, direct input from stimulated UCN3 to GABA efferent neurons will lead to the increase in its activity, consequently increasing KNDy inter-pulse interval (IPI). This finding contrasts with experimental observations, where the combined suppression of glutamate and stimulation of UCN3 failed to produce any significant change in GnRH pulsatility^23^.

Additionally, UCN3 neurons may co-release other fast neurotransmitters, such as glutamate^27^. The result of our UCN3 neurons stimulations at low frequencies (5 Hz; Supplementary Fig. 7), which primarily elicits the fast neurotransmitters release^25^, suggests that their isolated effects may be limited. We note that in our modeling framework, we include the constant drive, reflecting the effects of slow neuropeptide action, and a periodic drive corresponding to the action of fast neurotransmitters aligned with the stimulation frequency. The constant drive governs the overall dynamical regime, while the fast periodic term functions primarily as a rhythmic modulator, that is when the periodic drive dominates over the constant drive both the glutamatergic and GABAergic populations become entrained to the stimulation frequency. Nonetheless the specific contribution of co-transmitters remains experimentally unclear, and that may influence how the MePD integrates this tonic input (Fig. 7).

**Figure 7:**
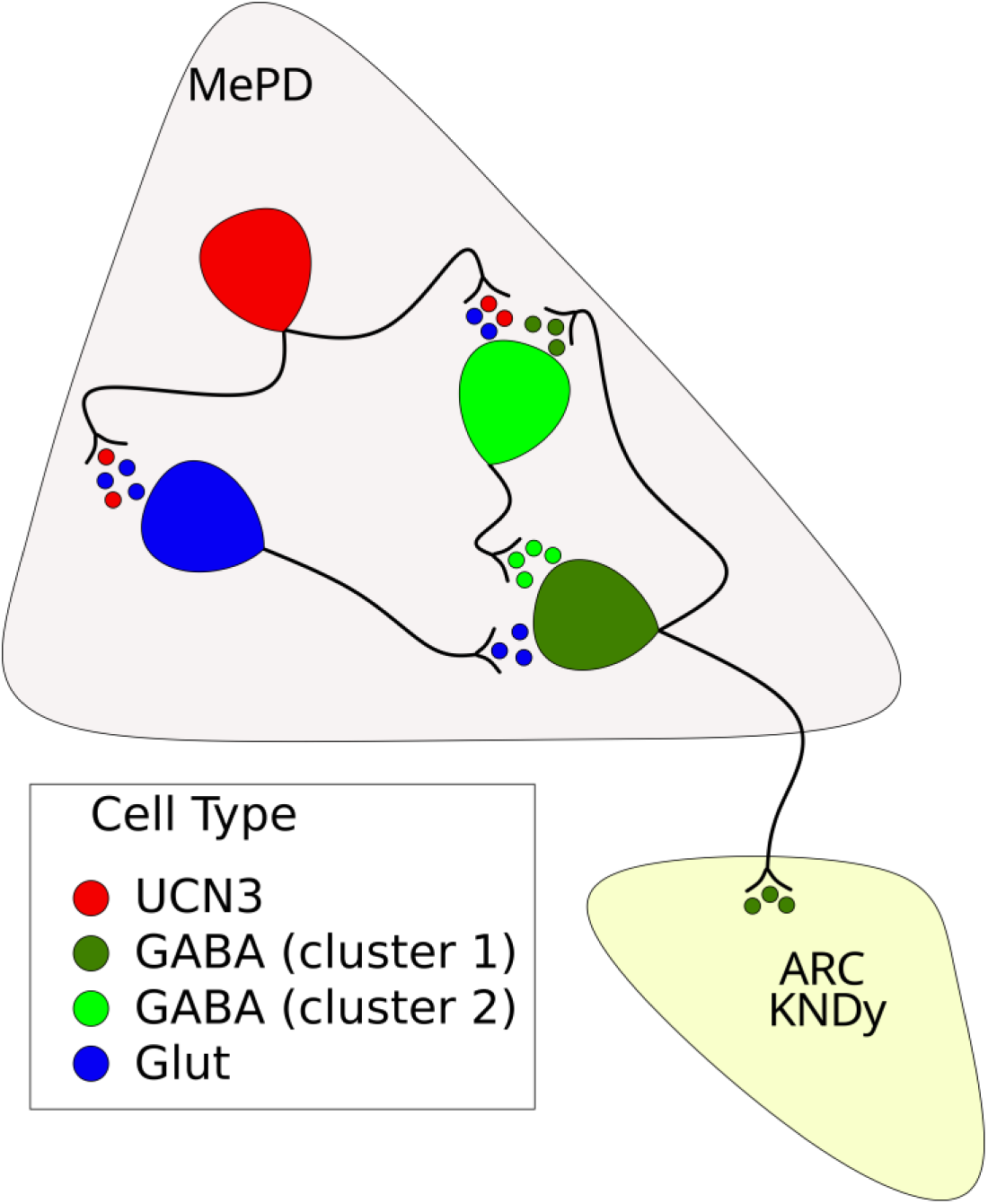
Schematic illustration of the proposed functional MePD neurocircuit. UCN3 neurons activate inhibitory GABA interneurons and excitatory glutamate neurons through the release of UCN3 and potentially other fast neurotransmitters. The glutamate and GABA interneuron populations then signal to the GABA efferent projection neurons, that convey the MePD output to the KNDy neurons in the arcuate nucleus.

We note that our MePD model has potential applications beyond understanding MePD dynamics and their influence on the GnRH pulse generator. The model incorporates two functionally distinct populations of GABA neurons (interneurons and efferent projection neurons), which reflect known differences in GABA neurons within the amygdala^18,20^. This diversity likely originates from the medial ganglionic eminence an embryonic structure that gives rise to GABA interneuron populations across various brain regions, including the MePD, through migration during embryogenesis^29^. Therefore, our modelling framework could be extended to other neural circuits where two functionally distinct GABA populations exist, such as the posterior ventral medial amygdala^20^ and its lateral and basal subdivisions^30^.

Consistent with model predictions, we showed that optogenetic activation of non-GABA neurons, expressing UCN3, suppresses LH pulse frequency. This finding corroborates previous work demonstrating that LH pulses are inhibited by activation of UCN3 neurons^13,23^ or administration of UCN3^16^ in the MePD, and that such effects are attenuated by chemogenetic inhibition of UCN3 neurons or by antagonizing corticotropin-releasing factor type 2 receptors (CRFR2)^16^. Thus, our findings further support the central role of UCN3 signaling in modulating both stress responses and reproductive function. Future studies could investigate the contribution of UCN3 neurons to stress integration more directly, for example by silencing UCN3 neurons during stress exposure to determine whether GABAergic activation and the associated network clustering patterns are abolished.

GABA is one of the primary inhibitory neurotransmitters in the nervous system. Previous work has shown that MePD GABA neurons project to the ARC to regulate LH pulses and chemogenetic inhibition of MePD GABA neurons attenuates stress-induced suppression of LH pulses^17^. Here, we found that optogenetic activation of GABA neurons in the MePD leads to robust LH pulse suppression, reinforcing the pivotal role of amygdala GABAergic signaling in stress-related reproductive dysfunction. Antagonizing GABA receptors in the MePD can block UCN3-induced LH suppression^23^, suggesting that GABA neurons may function downstream of UCN3 signaling.

However, this pharmacological approach lacks cell-type specificity, leaving open the possibility that UCN3 neurons themselves co-expressing GABA receptors, potentially contributing to the observed effect. Here, by using an intersectional genetic strategy, we demonstrated that optogenetically inhibiting non-UCN3 GABA neurons can abolish the change in LH pulse intervals induced by optogenetic stimulation of UCN3 neurons. This finding supports the hypothesis that these GABA neurons are acting downstream of UCN3 signaling, mediating LH pulse suppression. We note that while in our current experimental protocol we could also be blocking GABAergic projections to the ARC KNDy population, which by itself could explain the absence of any effect from UCN3 stimulation, our modeling predicts that GABAergic interactions within the MePD are additionally crucial for relaying UCN3 signals to the KNDy network as also suggested by pharmacologically antagonizing GABAreceptors^23^.

It is not unusual for neurons to co-express and co-release neuropeptides and neurotransmitters^20^. Here, our statistical analysis (Supplementary Fig. 2 & Supplementary Table 1) of GABA calcium traces indicate that certain GABA neurons exhibit activity entrained to the stimulation frequency of UCN3 neurons, suggesting a possibility that UCN3 is co-released with GABA. Given that activating either UCN3 or GABA neurons independently suppresses LH pulses, one might predict that stimulating neurons co-expressing both UCN3 and GABA would similarly inhibit LH. However, we observed no significant changes in LH pulses upon optogenetic stimulation of neurons co-expressing UCN3 and GABA. Our results indicate that UCN3 and GABA are both essential for transmitting inhibitory signals to the hypothalamic GnRH pulse generator; however, this inhibitory influence relies on their action through separate, functionally distinct neuronal populations. That is activation of neurons that exclusively express either UCN3 or GABA suppresses the GnRH pulse generator, whereas activation of UCN3 neurons that also co-express GABA fails to produce this inhibitory effect. Within our modeling framework, UCN3/GABA co-expressing neurons might be considered part of the GABA interneuron population, primarily inhibiting GABA projection neurons rather than activating them. Consequently, they fail to relay the inhibitory UCN3 signal to the ARC. Further studies are necessary to define more precisely the distinct role of UCN3/GABA co-expressing neurons within the MePD circuitry.

By combining mini-endoscopic calcium imaging with computational modeling, we identified two GABA subpopulations with complementary but distinct roles in encoding stress signals and modulating LH pulsatility. Our experimental and computational results also stressed the functional orthogonality between UCN3 and GABA signaling in MePD, highlighting the complexity of the MePD’s contribution to stress-induced reproductive suppression. These results offer improved understanding of the neuronal circuits in the amygdala, a key hub for emotional processing, involved in regulating hypothalamic GnRH pulse generator frequency^9–11,16,17,23^ providing new insight into psychogenic stress-related disorders of fertility, such as functional hypothalamic amenorrhoea^6,7,31,32^, and the development of novel therapeutic approaches to improve reproductive health and well-being.

## Methods

### Animal

VGAT-Cre-tdTomato mice were genotyped using PCR to determine heterozygosity for VGAT-Cre (primers 5′-3′: common, 12785—CTTCGTCATCGGCGGCATCTG; wild-type reverse 12786—CAGGGCGATGTGGAATAGAAA; mutant reverse oIMR8292—CCAAAAGACGGCAATATGGT) and tdtomato (primers 5′-3′: wild-type forward oIMR9020—AAGGGAGCTGCAGTGGAGTA; wild-type reverse oIMR9021—CCGAAAATCTGTGGGAAGTC; mutant reverse WPRE oIMR9103—GGCATTAAAGCAGCGTATCC; mutant forward tdTomato oIMR9105—CTGTTCCTGTACGGCATGG). UCN3-Cre heterozygous mice (strain Tg(Ucn3-cre)KF43Gsat/ Mmucd, MMRRC GENSAT) were crossbred with VGAT-Flpo heterozygous mice (Jax stock #007909,B6.Cg-Slc32a1 2A-Flpo-D:Het) to obtain female double heterozygous UCN3-Cre::VGATFlpo mice (Slc32a1 2A-Flpo-D:Het|Ucn3-Cre:Het|Rosa26<tomato>WT); genotyped using PCR for the detection of heterozygosity (primers 5′-3′: UCN3 Forward: CGAAGTCCCTCTCACACCTGGTT; Cre Reverse: CGGCAAACGGAC-AGAAGCATT; Slc32a1 Mutant Forward: TGC ATC GCA TTG TCT GAG TAG; Slc32a1 Mutant Reverse: GAC AGC CGT GAA CAG AAG G). Female mice, aged 8 to 10 weeks, weighing between 20 g and 25 g, were individually housed in ventilated cages sealed with a HEPA-filter at 25 ± 1°C in a 12:12-hour light/dark cycle, lights on at 07:00 h. Vaginal smears were performed when mice are 6 weeks old between 10:00 - 12:00 h daily. Only mice showing regular estrous cycles were included. All animal procedures performed were approved by the King’s College London Animal Welfare and Ethical Review Body. Procedures were in accordance with UK Home Office regulations.

### Stereotaxic injection of adeno-associated virus construct and fiber-optic or mini-endoscope GRIN lens implantation

All surgical procedures were carried out under general anesthesia using ketamine (Vetalar, 100 mg/kg, i.p.; Pfizer, Sandwich, UK) and xylazine (Rompun, 10 mg/kg, i.p.; Bayer, Leverkusen, Germany) under aseptic conditions. Animals were secured in a David Kopf stereotaxic frame (Model 900, Kopf Instruments, CA, USA). Following bilateral ovariectomy, a midline incision was made in the scalp to reveal the skull. Two small bone screws were inserted into the skull. A small hole was drilled above the position of the MePD. The MePD coordinate (2.29 mm lateral, 1.60 mm posterior from bregma, at a depth of 5.33 mm below the skull surface) was determined according to the mouse brain atlas of Paxinos and Franklin^33^. A unilateral stereotaxic viral injection was performed to target opsin or fluorescent protein expression in the MePD neurons, using a robot stereotaxic system (Neurostar, Tubingen, Germany), with a 5-μL Hamilton microsyringe (Esslab, Essex, UK), at the speed of 25 nl/min for 10 min.

Two transgenic animal models were used for LH blood sampling under optogenetic stimulation: UCN3Cre::VGAT-Flpo and VGAT-Cre-tdTomato. The following types of viral constructs were used in separated cohorts of UCN3-Cre::VGAT-Flpo mice: (i) AAV-nEF-Con/Foff 2.0-ChRmine-oScarlet (250 nl, 2.1×10^12^ gc/mL, Serotype:8, #137161, Addgene, Massachusetts, USA); (ii) AAV-nEF-Con/Fon 2.0ChRmine-oScarlet (250 nl, 1.9×10^12^ gc/mL, Serotype:8, # 137159, Addgene, Massachusetts, USA); (iii) a mixture of AAV-nEF-Con/Foff 2.0-ChRmine-oScarlet (2.1×10^12^ gc/mL, #137161) and pAAV-nEF-Coff/Fon-NpHR3.3-EYFP (1.9×10^12^ gc/mL; #137159, final volume 300 nl); (iv) control virus AAV-Ef1a-DIO-EYFP (250 nl, 4.4 × 10^12^ gc/mL, Serotype:9, #27056, Addgene, Massachusetts, USA). VGATCre-tdTomato mice received unilateral injections of (i) channelrhodopsin (ChR2) viral construct, (250 nl, 1.8 × 10^12^ gc/mL; Serotype:9; Addgene, #20298, Massachusetts, USA) or (ii) control virus, AAVEf1a-DIO-EYFP (250 nl, 4.4 × 10^12^ gc/mL, Serotype:9; Addgene, # 27056, Massachusetts, USA) into the MePD. Following injection, the needle was kept in position for 10 min before going up slowly over 1 min. A unilateral fiber optic cannula (200 µm, 0.39 NA, 1.25 mm ceramic ferrule; Doric Lenses, Quebec, Canada) was inserted into the right MePD 0.15 mm above the injection site. After reaching the target, the fiber optic cannula was secured on the skull using dental cement (Super-Bond Universal Kit, Prestige Dental, UK), bonded to the screws and skull. Finally, the incision in the scalp was closed with suture. Animals were given 1 week to recover, after which they were handled 20 min daily to acclimate to experimental procedures for a further 2 weeks.

For mini-endoscope *in vivo* calcium imaging, only UCN3-Cre::VGAT-Flpo transgenic animals were used. Separate mice cohorts received the following viral injection into the MePD: (i) a mixture of pAAV-Syn-Flex-rc[Chrimson-tdTomato] (2.0×10^12^ gc/mL, Serotype:5, #62723, Addgene, Massachusetts, USA) + pAAV-EF1a-fDIO-GCaMP6s (2.3×10^12^ gc/mL; Serotype:8; #105715, Addgene, Massachusetts, USA; final volume 400 nl); (ii) a mixture of pAAV-Syn-Flex-rc[ChrimsontdTomato] (2.0×10^12^ gc/mL, Serotype:5, #62723, Addgene, Massachusetts, USA) + pAAV.CAG.Flex.GCaMP6s.WPRE.SV40 (4.3×10^12^ gc/mL; Serotype:9; # 100842, Addgene, Massachusetts, USA; final volume 400 nL); (iii) a mixture of control virus pAAV-hSyn-DIO-mCherry (2.3×10^12^ gc/mL, Serotype:9, #50459) + pAAV-EF1a-fDIO-GCaMP6s (2.3×10^12^ gc/mL, Serotype:8, #105715, final volume 400nl). One week after administering the virus, a 600-µm diameter, 7.2-mm long gradient index (GRIN) ProView™ Integrated Lens (Inscopix, PN:100-004554) was implanted at the same location as the viral injection. The surface of the skull was scored, using the tip of a surgical scalpel, and four screws were inserted into the skull. This was done to enhance the adhesion of dental cement to the skull, thereby reducing the interference of movement on *in vivo* recording. Before insertion of the GRIN lens, a 23 G needle was slowly lowered at a speed of 5.0 mm/min into the brain (2.42 mm lateral, 1.52 mm posterior from bregma, depth of 5.05 mm below the skull surface) to make a needle track to facilitate the insertion of the GRIN lens without causing brain distortion along its route. The GRIN lens was then inserted into the brain at a rate of 0.6 mm/min, using the same coordinates as the needle precutting. To secure the GRIN lens with integrated baseplate in position, we used a light curing adhesive (iBond Universal, Kulzer GmbH, Leipziger, Germany) along with a cured radiopaque hybrid dental cement (Gradia Direct Flo, GC corporation, Tokyo, Japan), bonding it to the head screws and skull. A GRIN lens baseplate dust cover (Inscopix) was rescued in place to protect the lens. After 4 weeks the mice were re-anaesthetized as described above and underwent bilateral ovariectomy.

### Serial blood sampling for LH measurement

The mouse’s tail-tip was excised using a sterile scalpel. Then the chronically unilaterally implanted fiber optic cannula was connected to a multimode fiber optic rotary joint patch cables (Thorlabs Ltd, Ely, UK) via a ceramic mating sleeve to allow the animal freedom of movement. Mice were left for 1 h to habituate. A Grass SD9B stimulator-controlled DPSS laser (Laserglow Technologies, Toronto, Canada) was utilized to deliver either green light (532 nm wavelength, 10 Hz, 5 seconds on 5 seconds off, 10 mW, 10-ms pulse width,) or blue light (473 nm wavelength, 5 Hz, 5 seconds on 5 seconds off, 5 mW, 10-ms pulse width,). The light pattern was controlled using software designed in house. After 1 h of acclimatization, serial blood samples (5 µL) for LH pulse measurement were collected every 5 min for 2 h between 10:00 - 12:00 h: 0-60 min for control blood sampling period, and 60-120 min under stimulation or equivalent no-stimulation control period. Blood samples were diluted in 45 μL of Phosphate-Buffered Saline (PBS) containing 0.2% Bovine Serum Albumin (BSA) and 0.05% Tween20 (PBST) and immediately placed on dry ice and stored at −80°C until later analysis.

### LH assay

LH measurement was performed using an LH ELISA assay as reported previously^34^. A mouse LH standard (mLH; AFP-5306A, NIDDK-NHPP, USA) was used to establish a standard curve for the quantitative measurement. The following antibodies were used in the assay: (i) coating antibody (RRID: AB_2665514, monoclonal antibovine LH beta subunit antiserum, 518B7, University of California, CA, USA); (ii) anti-LH antibody (RRID: AB_2665533; National Hormone & Peptide Program, CA, USA); (iii) secondary antibody (RRID: AB_772206, GE Healthcare, Chicago, Illinois, USA). The inter-assay and intra-assay variations were 10.53% and 5.74%, respectively. The assay sensitivity was 0.023 ng/mL.

### LH pulse statistical analysis

LH pulses were verified using the Dynpeak algorithm in Scilab 5.5.2 programme^35^. The Dynpeak algorithms were classified for OVX mice based on previously established parameters^36^. The average LH inter-pulse interval was calculated for the 1-h control period and the 1-h stimulation period in both experimental and control mice. Statistical significance was assessed using a 2-way ANOVA unless otherwise stated. Data were represented as mean ± SEM and p<0.05 was considered significant.

### Mini-endoscope-based Calcium imaging

**Optogenetic stimulation of MePD UCN3 neurons.** Prior to recording, animals were handled daily for 5 to 6 weeks to get accustomed to the experimental setup. The Data Acquisition (DAQ) box (Inscopix, PN:100-005904) was turned on and connected to the computer wirelessly. The Data Acquisition Software (IDAS; Inscopix, Inc.) was accessed through the Chrome web browser on the computer communicating with the DAQ box. Adequate DAQ box storage capacity was checked routinely before Calcium imaging. The GRIN lens baseplate dust cover was gently removed from the mouse’s head, after which the minscope camera was secured to the baseplate platform. Equipped with the head-mount miniscope (nVoke2, Inscopix, SN: BB-11238601), the mouse was then placed back into its home cage. The optimal focal plane was chosen by adjusting the lens focus to achieve clear cell morphology. The exposure time was fixed at 50 ms. The frame rate was set to 20 Hz. Subsequently, gain, and LED power were optimized based on image histograms before initiating recording. Two types of LED were used during the recording: (i) EX-LED (0.2-0.4 mW, 435-460 nm excitation filter) for GCaMP6s signal detection in the GABA or UCN3 neurons; (ii) OG-LED (10 mW, 590-650 nm excitation filter) for Chrimson optogenetic stimulation of the UCN3 neurons. After the mice were attached to the miniscope, they were given 1-2 h for adaptation. For stimulation of the UCN3 neurons, the OG-LED was set to 10 Hz, 5 sec on 5 sec off, and a 10-ms pulse width. Once the animals had acclimatized, the GCaMP6s recording of the GABA neurons or UCN3 neurons was started with a protocol of 2 min baseline before stimulation for 2 min. Optogenetic stimulation was randomized for the 10 Hz, with a 5-min interval between each to prevent photobleaching of GCaMP6s. Once the experiment was concluded, the miniscope was gently detached, and the baseplate cover was reattached and secured with its screw. The recordings were performed between 9:00 h and 15:00 h. Three groups of animals were included in this experiment: (i) mice with UCN3 neurons expressing opsin and GABA neurons expressing GCaMP; (ii) mice with UCN3 neurons expressing both opsin and GCaMP, serving as a positive control; (iii) mice with UCN3 neurons expressing mCherry and GABA neurons expressing GCaMP, serving as a negative control. To quantify intra-animal variability at least two independent stimulation trials were conducted with each animal, with a maximum of four trials per animal.

**Restraint stress.** The system was set up and the mouse was connected to the miniscope as described above but with the OG-LED turned off. Once the mouse had settled, the experiment started with a 3 min baseline recording, followed by 3 min of recording under hand-held restraint. Finally, the miniscope was detached and the baseplate cover was reattached. The experiments were performed between 9:00 h and 15:00 h. The animals in this experiment were the same as those used in the optogenetic stimulation protocol described above but were divided into two groups depending on whether GCaMP was expressed in the UCN3 or GABA neurons, regardless of whether opsin or mCherry was expressed in the UCN3 cells. A single trial per animal was performed to minimize unnecessary distress, in accordance with the 3Rs principle, apart from one animal in which the restraint stress experiment was performed twice to ensure this does not result in habituation.

### *In vivo* calcium imaging recording processing

The video data, with a resolution of 1280 pixels x 800 pixels, was exported from the DAQ box through the file manager to a desktop computer and processed with the Inscopix Data Processing Software (IDPS). Within the IDPS, there are six essential steps for extracting single-cell calcium transient data from the recorded video: down-sampling, cropping, spatial filtering, motion correction, trace normalization (ΔF/F_0_), and cell identification. First, the video underwent both spatial down-sampling and temporal down-sampling by a factor of 2 and was cropped to enhance processing speed, focusing on the region of interest where the GCaMP signal was observed. Spatial filtering, with a threshold ranging between 0.005 (low) and 0.5 (high) was used to distinguish individual pixels in the video image, creating a high-contrast image. A mean intensity projection image was generated. Motion correction was subsequently conducted, using the mean intensity projection image as a reference. Motion correction was applied again using one of the frames affected by movement while simultaneously drawing a region of interest (ROI) where UCN3 accumulates to enhance the effectiveness of motion correction. For the recordings with GCaMP signals from GABA neurons, we did not use the built-in motion correction function of IDPS. Instead, we used the MATLAB (version R2022a) implementation of the open-source algorithm NoRMCorre^37^ with default parameters for both rigid and non-rigid (patch size 64 pixels and overlap 32) motion correction as long as a minimum correlation of the frames with the mean frame above 0.8 is achieved. This was performed by doubling the patch size and their overlap to 128 and 64, respectively. Corrected videos were saved in TIFF format for further analysis via IDPS. Following motion correction, pixel intensity, *F*(*x*, *y*), in each frame of the movie was normalized using the following formula 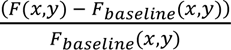, where *F*_*baseline*_(*x*,*y*) the baseline pixel intensity calculated by taking the mean value of the pixel across the entire movie. Finally, regions of interest were identified automatically using the cell identification algorithm provided by IDPS, based on principal component and independent component analysis (PCA-ICA algorithm).

### Validation of virus injection, fiber optic cannula and GRIN lens implantation

Once the experimental procedures were completed, a lethal dose of ketamine was used to humanely euthanize the mice. The mice were perfused transcardially with heparinized saline (5 U/ml) for 5 min, followed by ice-cold 4% paraformaldehyde (PFA) for 15 min using a pump (Minipuls,156 Gilson, Villiers Le Bel, France). The brains were collected and post-fixed sequentially at 4°C in a solution of 15% sucrose in 4% PFA and later in a solution of 30% sucrose in phosphate buffer until they sank. They were then snap frozen on dry ice and stored in −80°C until processing. The brains were coronally sectioned (30 μm) between +0.5 mm and −2.7 mm from the bregma using a cryostat (Bright Instrument Co., Huntingdon, UK), and sections were mounted on microscope slides, air-dried, and covered with slips using Prolonged Antifade mounting medium (Molecular Probes, Inc. OR, USA). The accuracy of injection sites, fiber optic cannula and GRIN lens implantation were confirmed by Axioskop 2 Plus microscope equipped with AXIOVISION 4.7 (Zeiss). Images were captured using Axioskop 2 Plus microscope (Carl Zeiss). Only the data of mice with correct viral infection and implantation of cannula and lens in the MePD were included in the analysis.

### Immunohistochemistry

To assess the targeting of viral labeling to UCN3 neurons in the MePD, we used the indirect immunofluorescence technique to localize UCN3 neurons. Every 4^th^ section from each brain was washed in potassium phosphate-buffered saline (KPBS) and incubated in blocking solution (KPBS containing 1% BSA, 0.03% Triton X-100 and 5 mg/mL heparin) at room temperature for 1 hour followed by incubated in a rabbit polyclonal anti-UCN3 antibody (1:2000 dilution)^38^ in the blocking solution (Code: PBL 7218, kindly provided by J. Vaughan, Salk Institute, La Jolla, CA, USA) for 48 hours at 4°C. The brain sections were then processed with goat anti-rabbit IgG (H + L), a biotinylated secondary antibody (1:1000 dilution; Cat. No. BA-1000, Vector Laboratories, Burlingame, CA, USA) overnight at 4°C. After rinses in KPBS, visualization of UCN3 immunoactivity was achieved using Streptavidin, Alexa Fluor 405 conjugate (1:200 dilution; Cat. No. S 32351, Thermo Fisher Scientific, Waltham, MA, USA) for 2 hours at room temperature. The sections were finally washed, mounted onto slides and cover slipped. Omission of the UCN primary antibody and its pre-incubation resulted in the absence of specific staining. Brain sections from each experimental group were processed on the same day to control for inter-batch variability. To preserve the fluorescence of the virally expressed proteins, all subsequent steps were performed in a dark box. Semiquantitative analysis of immunostaining data was carried out on AxioVision microscope image system. UCN3-positive neurons were identified by Alexa Fluor 405 fluorescence, while native fluorescence (e.g. tdTomato or oScarlet) was used for virus-transduced neurons identification. The number of virus-labeled, UCN3-immunoreactive cells and co-localized cells in the MePD were quantified on 4 sections from each mouse brain. Targeting efficiency was defined as the percentage of virus-labeled cells that were also positive for UCN3.

### Data analysis

The data processing protocol we employ in this study was performed in two phases: pre-intervention and intervention (either via optogenetic stimulation or restraint stress). For our analysis, we use the smoothed deconvolved traces generated by the OASIS method^39^, which improves the signal-to-noise ratio (SNR) and mitigates noise interference. All traces are rescaled between 0 and 1 (min-max normalization), and then further normalized to mean zero and unit variance. The analysis process is divided into three key steps: i) identification of GABA cells co-expressing UCN3 and removing them from further analysis; ii) estimating and comparing functional connectivity before and during intervention; iii) clustering analysis based on pairwise association-derived dissimilarity scores. In step (i), we use power spectral density (PSD) analysis and K-means clustering (using Euclidean distance) to identify GABA neurons with activity that mimics the pattern of optic stimulation, evidence that they are co-expressing UCN3.

**PSD filtering.** For each trace, we compute the PSD during the stimulation phase. Cells with PSD values at 0.1 Hz (the frequency of the stimulus 5 sec and 5 sec off cycle) above 0.37 Hz^−1^ are labelled as co-expressing UCN3. This threshold is determined from recordings of UCN3 cells under optic stimulation, by calculating the median of the minimum PSD values at 0.1 Hz from each dataset (see Supplementary Fig. 2**B**).

**K-means clustering.** Using the Euclidean distance metric, we cluster the traces during the stimulation phase into two groups, aiming to separate neurons into GABA-only neurons and neurons co-expressing UCN3 and GABA. Clustering separated a small group of neurons with activity mimicking the optic stimulation oscillatory pattern (see Supplementary Fig. 2**C**). Recordings of MePD UCN3 neurons shows that under optogenetic stimulation their activity rises and declines in phase with on and off periods of stimulation (shown in Fig. 3**A**); therefore, this small number of cells was labeled as co-expressing UCN3.

The union of neurons identified by either method as co-expressing UCN3 and GABA are excluded from further analysis. By applying both filters, we aim to robustly identify purely GABA-expressing neurons. We acknowledge the possibility that some cells may pass through the filters or that a few purely GABA-expressing neurons may be inadvertently excluded. However, our dual approach of employing PSD analysis and K-means clustering minimizes this possibility and is essential to ensure our analysis remains focused exclusively on GABA-expressing neurons. Nevertheless, we take a further step and visually inspect the calcium recording traces for any potential anomalies such as regular oscillations in-phase with the stimulation and/or massive increase in calcium level during the intervention period. Any identified anomalies of this sort are also dropped from datasets.

### Comparing pre-intervention and intervention cell-cell interaction strength

Following step (i), the remaining dataset contains only traces associated with GABA neurons, which are not directly activated by our optic stimulation protocol. In step (ii), we assess the functional connectivity between the observed neurons using Pearson’s correlation^40^ defined as

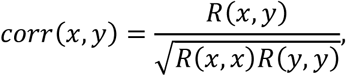

where 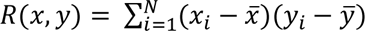 represents cross-correlation with 𝑁 as the length of 𝑥 and 𝑦. Using the above measures, we construct the corresponding cross-correlation matrices (see for an example Fig. 1**H**&**I**).

To evaluate changes in pairwise cell interactions between the pre-intervention and intervention phases, we compute the pairwise Riemannian distance^41^ between correlation matrices from pre-intervention and intervention episodes. The Riemannian distance here is defined as

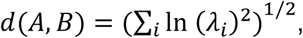

where 𝑙𝑛 is the natural logarithm and 𝜆. indicates the joint eigenvalues of symmetric positive definite (SPD) cross-correlation matrices 𝐴 and 𝐵. The size of the matrices directly affects the Riemannian distance, therefore, we normalize each Riemannian distance by the average Riemannian distance computed from 100 pairs of randomly generated SPD matrices of the same size. To make sure that random generation is uniform in the space of SPD matrices, we employ the Metropolis-Hasting method introduced in^42^. Normalizing the Riemannian distances guarantees that the distance is not influenced by matrix size. To assess the effect of experimental conditions on Riemannian distances while accounting for the hierarchical structure of the data, we employed a linear mixed-effects model^43^. Specifically, we modeled repeated measurements nested within individual animals, as follows:

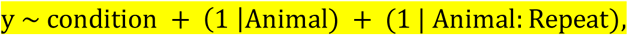

where 𝑦 denotes the computed Riemannian distance, while condition represents whether the experiment was control, stimulation or restraint stress. The model was fitted using maximum likelihood estimation.

### Clustering analysis

In step (iii), we apply agglomerative hierarchical clustering (using the average linkage criterion; package in scikit-learn Python™) to classify cells based on the dissimilarity of their calcium activity, given as

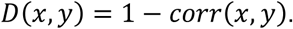

To determine the optimal number of clusters and robustness of clustering, we employed a bootstrapping technique. This involved resampling (without replacement) 75% of the cells and clustering the resulting dataset with different numbers of clusters (2, 3, and 4) using hierarchical or K-means clustering with either the covariance or correlation as the dissimilarity measure. This resampling process was repeated 100 times.

For each resample dataset, we evaluate the clustering performance, using the Silhouette Coefficient (SC)^42^ which measures the cohesion within clusters and their separation, with values ranging from −1 (poor performance) to 1 (good performance). Results of the evaluation are given in Supplementary Figs. 4-6, showing that the SC is significantly higher for 2 clusters compared to 3 or 4 clusters in the case of both hierarchical and K-means clustering.

### Modeling

We build on our previously published modeling framework of MePD circuit^11^ by incorporating dynamic stimulation by UCN3 neurons as observed experimentally (as shown in Fig. 3). To this end we define input from UCN3 neurons into the populations of glutamatergic neurons and GABA interneurons with a strength given by the parameter 𝛾. The activity of UCN3 neurons during stimulation is approximated via sinusoidal function:

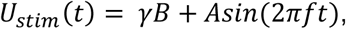

where the parameter 𝑓 is the frequency of the wave. Parameter 𝐵 represents the constant excitatory drive, potentially mediated by slow neuropeptide action, while the term 𝐴𝑠𝑖𝑛(2𝜋𝑓𝑡) describes faster neurotransmitter-driven-driven input aligned with optogenetic stimulation frequency. We approximate UCN3 neuron activity during restraint stress using two exponential functions:

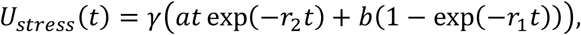

where the function initially increases with magnitude 𝑎 then decreases to a plateau value given by the parameter 𝑏. The parameters 𝑟_2_ and 𝑟_1_ are responsible for the rate of increase of the first term and rate of decrease to a plateau in the second term.

Briefly, the model captures the dynamics of the MePD intracellular calcium activity of three populations: glutamatergic neuronal population (excitatory) and the populations of GABA interneurons and GABA efferent neurons (inhibitory). We build on our previous work^11^ by assuming that excitatory (UCN3) input is provided to the populations of GABA interneurons and glutamate (Fig 3). Previous analysis of the model indicated that negative feedback between GABA and glutamatergic neuronal populations as well as glutamatergic self-excitation are critical mechanisms for inducing oscillatory dynamics^11^. However, providing glutamatergic inputs to both populations of GABA neurons results in a null phase difference between the populations. Therefore, we opted to provide glutamatergic input only to the population of GABA efferent neurons.

To investigate how the amygdala modulates the activity in the GnRH pulse generator, we couple the above-described MePD model to our KNDy network model, where the oscillatory dynamics of the KNDy acts as a proxy of LH pulsatility^26^. Here we couple the model via providing only inhibitory output from MePD GABA efferent neurons population to KNDy, as it was previously shown that glutamate plays a modulatory role to the MePD output rather than directly affecting the KNDy network^11^. We then calibrate the coupled model by reproducing the experimental results from Ivanova et al.^23^ (Supplementary Fig. 8). Further modeling details, such as equations governing model dynamics and model parameter values, can be found in Supplementary Information. The simulations have been done in MATLAB (version R2023b).

## Data Availability

Source data are provided with this paper. Further information that supports the findings of this study is available from the corresponding authors upon request.

## Code Availability

The code for reproducing the data analysis (Python™), mathematical modeling (MATLAB), and figure generation for both the main text and Supplementary information, along with the datasets used in the analysis, is publicly available at: https://github.com/mv-kr/MePD_GABA.

## Acknowledgements

We thank Ms Sumi Mathew for her technical support with genotyping mice.

S.F., M.V., K.T.A., K.T.O., X.F.L. gratefully acknowledge the financial support of BBSRC via grants BB/W005913/1 (KCL), BB/W005883/1 (Exeter) and BB/S019979/1. KTA gratefully acknowledges the financial support of the EPSRC via grant EP/T017856/.

## Contributions

J.Y. contributed data acquisition and data analysis, S.F. contributed data analysis, K.N. contributed modeling, M.V. contributed managing data analysis, X.F.L. contributed managing data acquisition and analysis, H.Y. contributed data acquisition, Y.L. contributed data acquisition, J.Y. contributed data acquisition, O.H. contributed data acquisition, R.D.B. contributed data acquisition, B.S. contributed data acquisition, K.T.O., K.T.A., X.F.L. and M.V. contributed to managing the project. All authors contributed to writing the manuscript.

## Competing interests

The authors declare no competing interests.

## Open Access

For the purpose of open access, the authors have applied a ‘Creative Commons Attribution (CC BY) licence to any Author Accepted Manuscript version arising from this submission.

## Notes

### Competing Interest Statement

The authors have declared no competing interest.

### Summary of Updates

Updated figure 3, added figure 7, expanded discussion

