## Supplementary Information for "GABA Neurons in the Amygdala play a central Role in Urocortin-3–Mediated Stress Suppression of Reproduction"

**[1]Department of Women and Children's Health, School of Life Course and Population Sciences, King's College London, Guy's Campus, London SE1 1UL, UK, [2]Department of Rehabilitation Medicine, The First Affiliated Hospital of Wenzhou Medical University, Wenzhou, Zhejiang 325000, China, [3]Department of Mathematics and Statistics, Stocker Road, Exeter EX4 4PY, UK, [4]Living Systems Institute, Exeter EX4 4QD, UK, [5]The Pirbright Institute, Ash Road, Pirbright, Surrey GU24 0NF, UK, [6]Biological Sciences, University of Missouri, Columbia, MO 65211-7400 and [7]EPSRC Hub for Quantitative Modelling in Healthcare, University of Exeter, Stocker Road, Exeter EX4 4PY, UK**

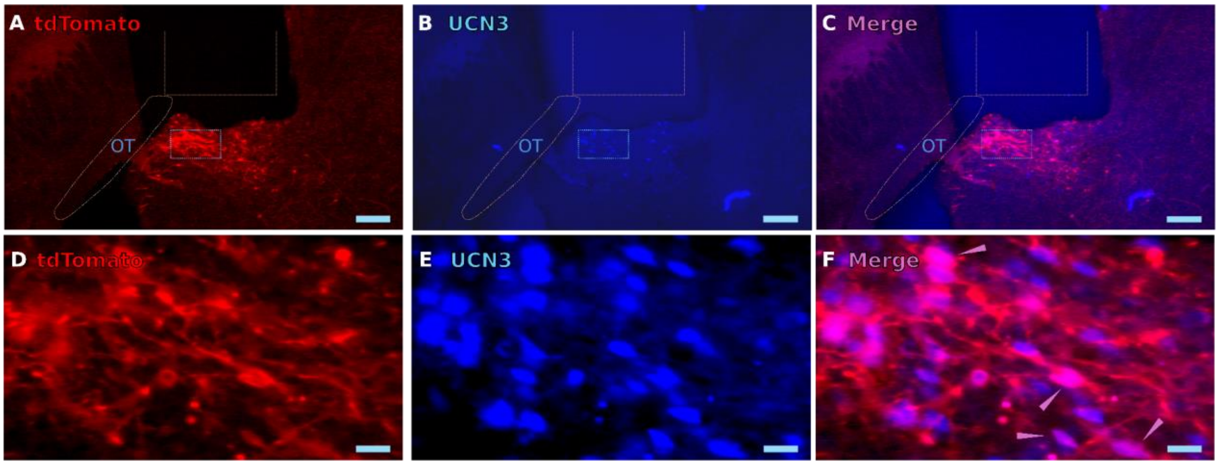


**Supplementary Fig. 1: Validation of Cre-dependent Chrimson-tdTomato expression in UCN3 neurons in the MePD.** (**A**&**D**) Red fluorescent labelled UCN3 neurons expressing Chrimson-tdTomato. (**B**&**E**) Blue fluorescent labelled UCN3 neurons immunopositive for the UCN3 antibody tagged with Alexa Fluor 405. (**C**&**F**) The merged images (magenta) demonstrate the co-localization of Cre-dependent Chrimson-tdTomato and UCN3 immunoreactivity. Pink arrowheads indicate double-labeled neurons. Position of the GRIN-lens is indicated. Scale bars represent (**A** to **C**) 200 μm, (**D** to **F**) 25 μm. OT, optic track.

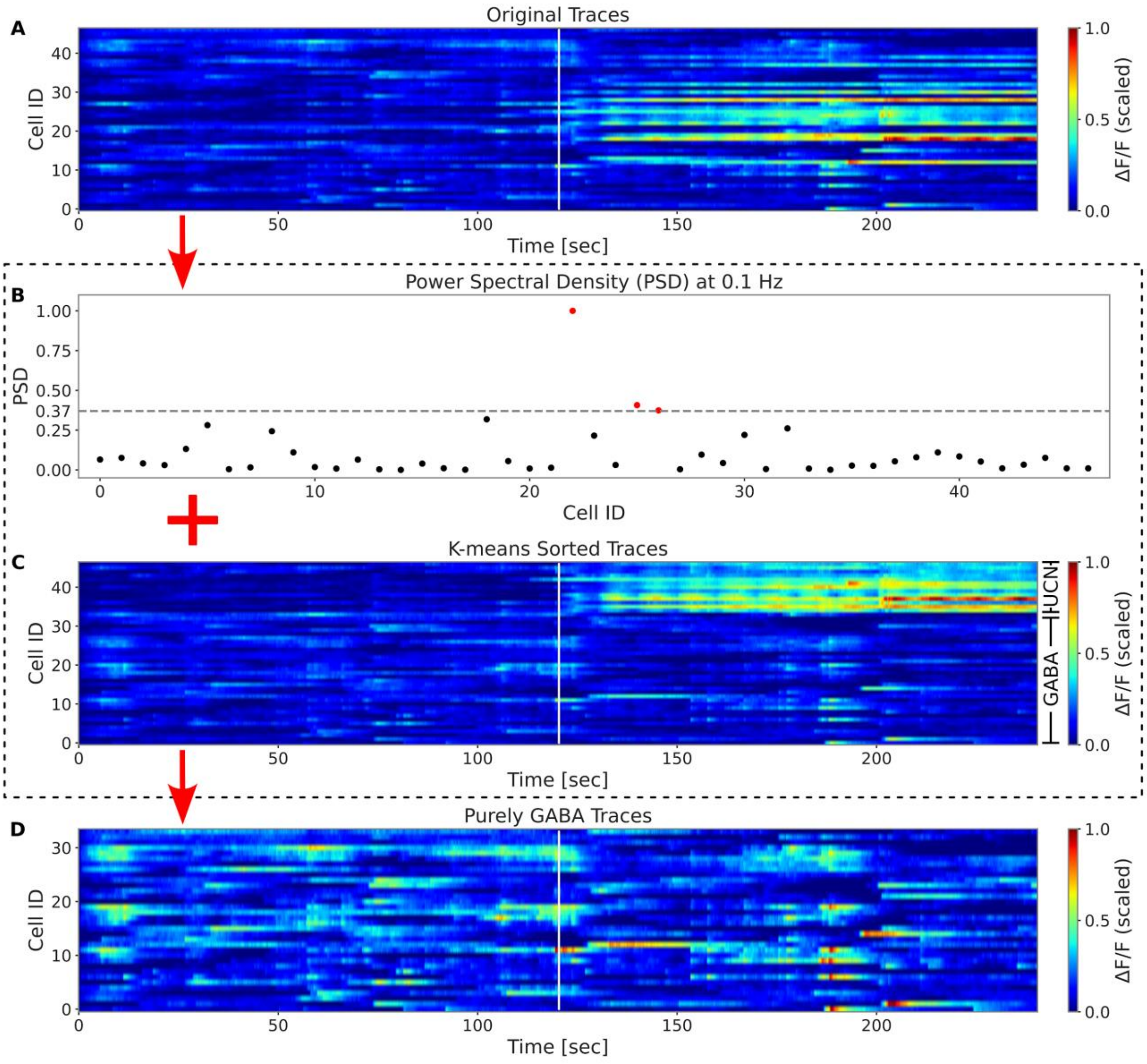


**Supplementary Fig. 2: Procedure for identifying and filtering neurons entrained to the UCN3 optogenetic stimulation frequency.** (**A**) The procedure is demonstrated on the dataset presented in Fig. 1 of the main text. Traces of 47 GABAergic neurons recorded during an optogenetic stimulation trial. (**B&C**) The procedure utilizes two distinct filters, one based on power spectral density (PSD) and the other on K-means clustering, to identify GABAergic cells that co-express UCN3. (**B**) Cells with PSD values at 0.1 Hz (the frequency of stimulation) exceeding 0.37 $Hz^{-1}$ are identified as co-expressing UCN3 and are indicated as red dots in the plot. A set of independent optogenetic stimulation trials where MePD UCN3 neurons were being recorded, was used to derive the threshold value as the median (across trials) of the minimum (over cells in each trail) PSD value at 0.1 Hz. (**C**) Cells co-expressing UCN3 appear as a small group (labeled

UCN) after K-means clustering while purely GABA cells comprise a bigger second cluster (labeled GABA). In total, 13 cells were identified as UCN3 cells using both filters. (**D**) Traces of the remaining 34 GABA neurons.

**Supplementary Table 1: Number of recorded cells, cells entrained to the UCN3 stimulation frequency, and final number of GABA cells after exclusion procedure.**

| Animal | # of detected cells | # of cells entrained to the UCN3 stimulation frequency | # of GABA cells post exclusion procedure |
|---|---|---|---|
| G7 | 60 | 8 | 52 |
| G7 | 47 | 13 | 34 |
| G8 | 65 | 10 | 55 |
| G8 | 42 | 14 | 28 |
| G8 | 56 | 23 | 33 |
| G8 | 69 | 23 | 46 |
| G9 | 46 | 19 | 27 |
| G9 | 44 | 17 | 27 |
| G9 | 51 | 14 | 37 |
| G9 | 48 | 17 | 31 |
| G28 | 38 | 9 | 29 |
| G28 | 41 | 5 | 36 |
| G28 | 47 | 4 | 43 |
| G29 | 31 | 18 | 13 |
| G29 | 31 | 24 | 7 |
| G29 | 30 | 18 | 12 |
| G37 | 40 | 18 | 22 |
| G37 | 36 | 21 | 15 |
| G37 | 38 | 15 | 23 |
| G37 | 32 | 17 | 15 |

**Supplementary Table 2: Number of recorded cells during the restraint procedure.**

| Animal | # of detected cells |
|---|---|
| G7 | 74 |
| G8 | 79 |
| G9 | 63 |
| G28 | 35 |
| G29 | 29 |
| G37 | 39 |
| G37 | 31 |

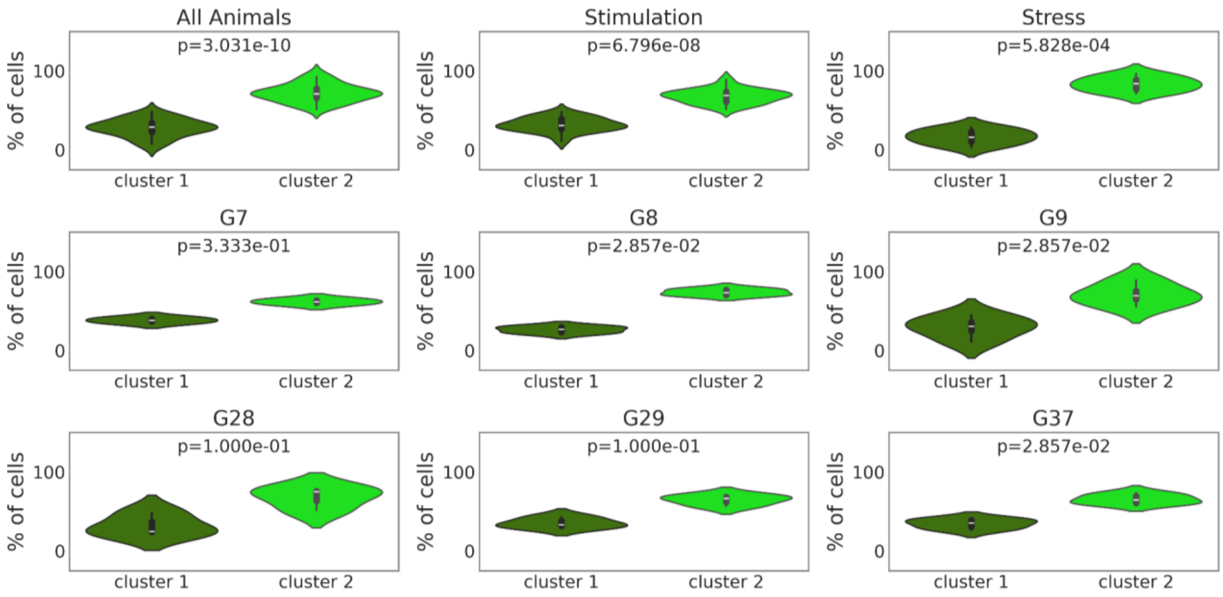


**Supplementary Fig. 3: Sizes of the two clusters observed in the GABAergic population under different conditions.** The percentages of GABAergic cells for each cluster in all animal trials, only in UCN3 optic stimulation trials, only in stress trials, as well as in all trials (both optogenetic stimulation and stress) per individual animal (G7-G37). The p-value of the statistical significance in size difference is shown at the top of each panel.

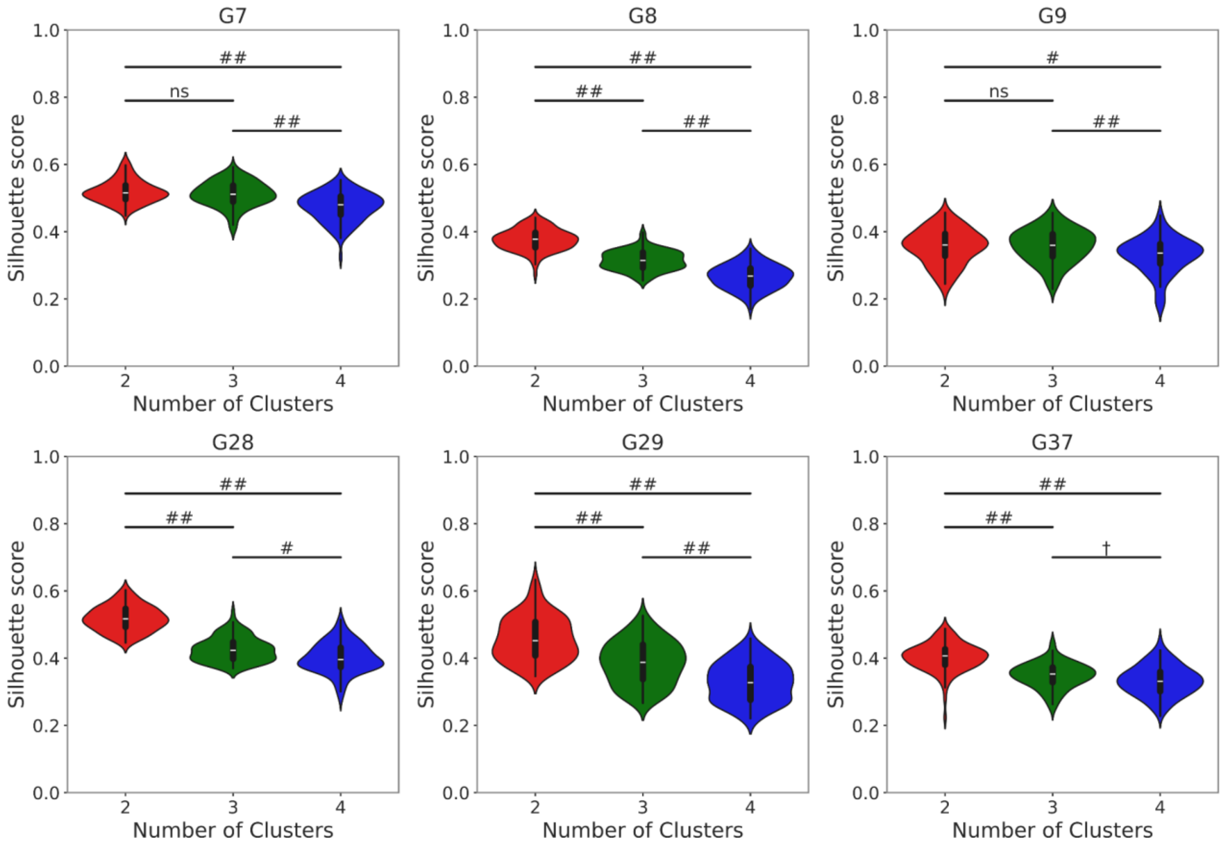


**Supplementary Fig. 4: The quality of hierarchical clustering using Pearson correlation-derived dissimilarity measure declines for more than two clusters**. Distribution of Silhouette scores obtained by resampling 75% of the GABA population, observed in a representative optogenetic stimulation trial for each animal, and using this resampled dataset to perform hierarchical clustering for different number of clusters (2, 3 and 4). The post-hoc Dunn statistical test with Benjamini–Hochberg correction method confirms that the hierarchical clustering method using Pearson correlation-derived dissimilarity measure robustly performs superior with 2 clusters compared with higher number of clusters. (†: $p<0.05$, #: $p<0.01$ and ##: $p<0.001$) Pearson correlation does not consider any delay between the calcium signals. In Supplementary Fig. 4, we use a modified cross-correlation, also accounting for lag and sign in cell-cell interactions, termed *signed lagged cross-correlation* (SLxCorr) which is adapted from the works of Tsuyuzaki et al.[1] and Paparrizos & Gravano[2]. It is defined as

$$SLXCorr(x,y) = \frac{R_{\tau_{max-N}}(x,y)}{\sqrt{R_0(x,x)R_0(y,y)}},$$

in which

$$\tau_{max} = \left( \left| \frac{R_{\tau-N}(x,y)}{\sqrt{R_0(x,x)R_0(y,y)}} \right| \right),$$

where $R_\tau = \{\sum_{i=1}^{N-\tau-1}(x_{i+\tau} - \bar{x}).(y_i - \bar{y})\ for\ \tau \geq 0\ R_{-\tau}(y,x)$ for $\tau < 0$ represents cross-correlation with lag $\tau \in \{-N+1, \ldots, -1, 0, 1, \ldots, N-1\}$ with $N$ as the length of $x$ and $y$. (##: p<0.001, #: p<0.01, †: p<0.05).

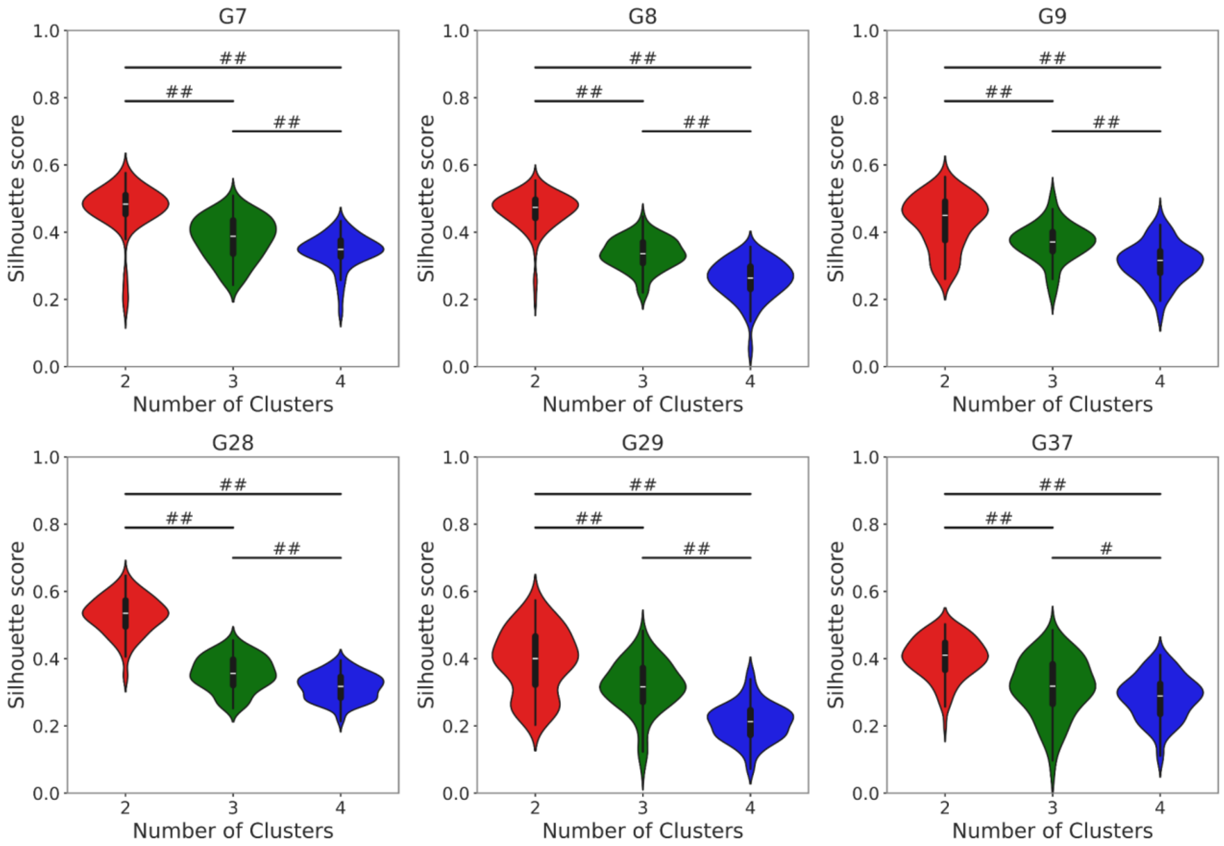


**Supplementary Fig. 5: The quality of hierarchical clustering using SLxCorr-derived dissimilarity measure declines for more than two clusters.** Distribution of Silhouette scores obtained by resampling 75% of the GABAergic population, observed in a representative optogenetic stimulation trial for each animal, and using this resampled dataset to perform hierarchical clustering for different number of clusters (2, 3 and 4). The post-hoc Dunn statistical test with Benjamini–Hochberg correction method confirms that the hierarchical clustering method using SLxCorr-derived dissimilarity measure robustly performs superior with 2 clusters compared with higher number of clusters. (##: p<0.001, #: p<0.01).

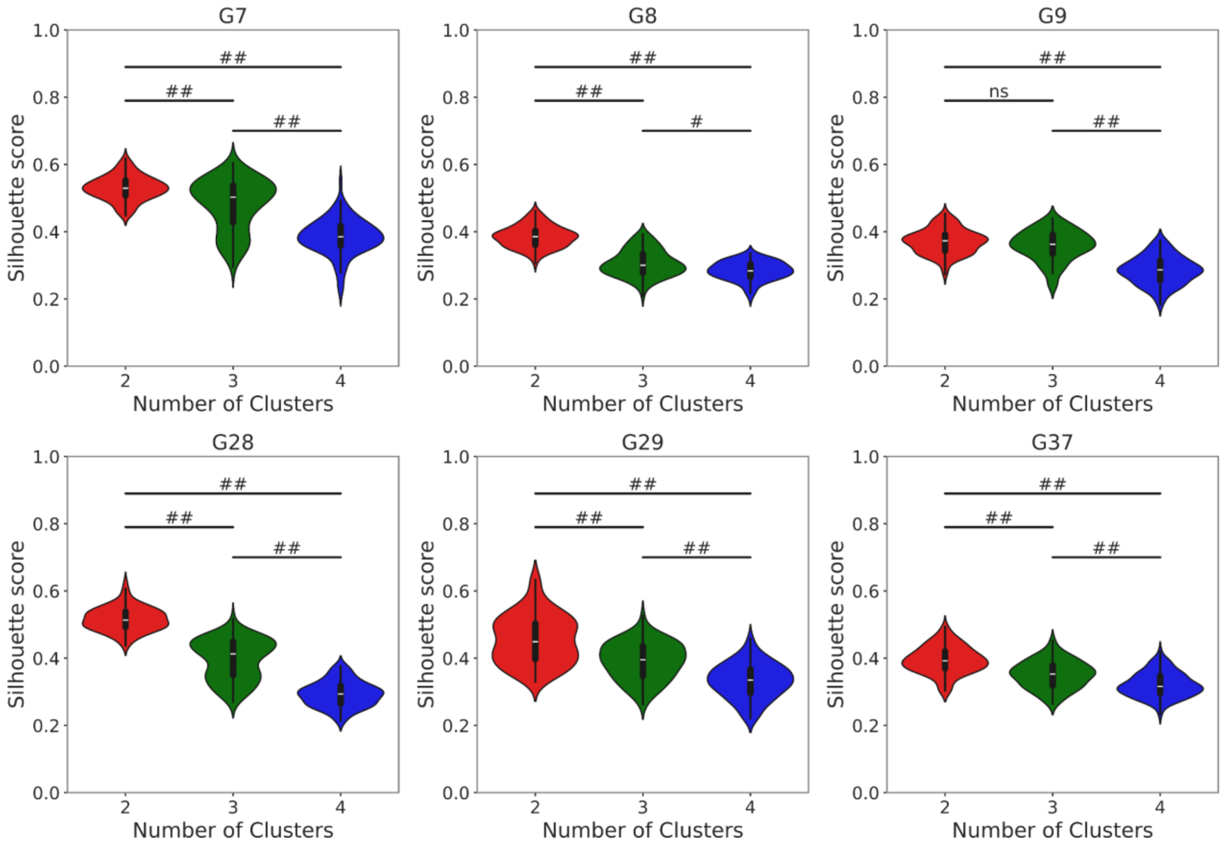


**Supplementary Fig. 6: The quality of K-means clustering using the Pearson correlation derived dissimilarity measure declines for more than two clusters.** Distribution of Silhouette scores obtained by resampling 75% of the GABAergic population, observed in a representative optogenetic stimulation trial for each animal, and using this resampled dataset to perform K-means clustering for different number of clusters (2, 3 and 4). The post-hoc Dunn statistical test with Benjamini–Hochberg correction method confirms that the hierarchical clustering method using Pearson correlation robustly performs superior with 2 clusters compared with higher number of clusters. (##: p<0.001).

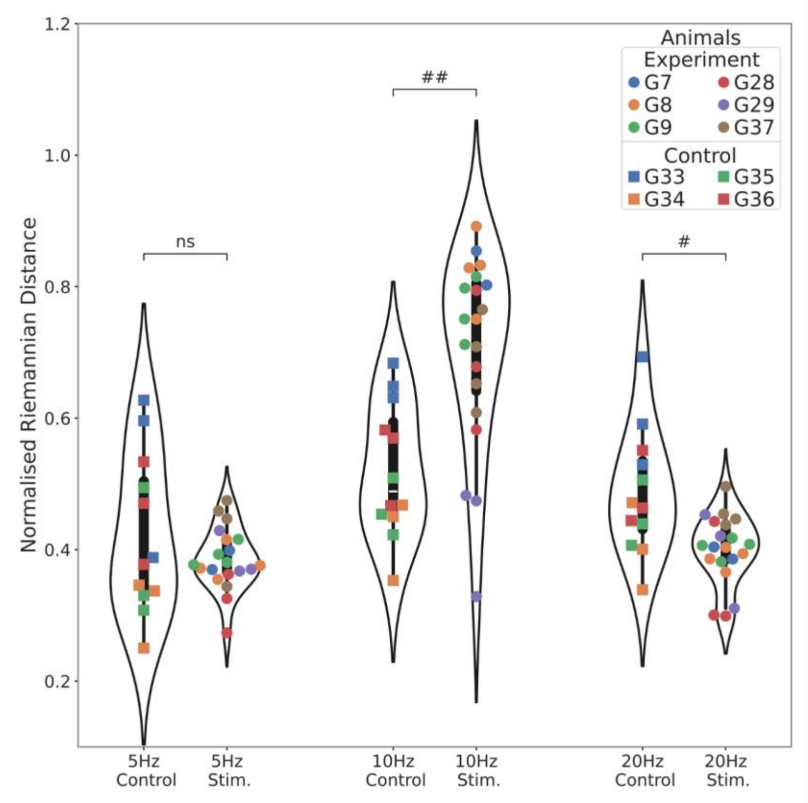


**Supplementary Fig. 7: Effect of stimulation strength/frequency on functional connectivity of the GABA network.** Applying a non-parametric Kruskal-Wallis statistical test shows that there are no significant changes for 5 Hz stimulation compared with control trials (left) while the effect of 10 Hz (middle) and 20 Hz (right) stimulations is significant. Note that 20 Hz stimulation gives rise to a decrease in Riemannian distance between Pearson correlation matrices of control and experimental trials. (##: $p<0.001$, #: $p<0.01$, ns: non-significant).

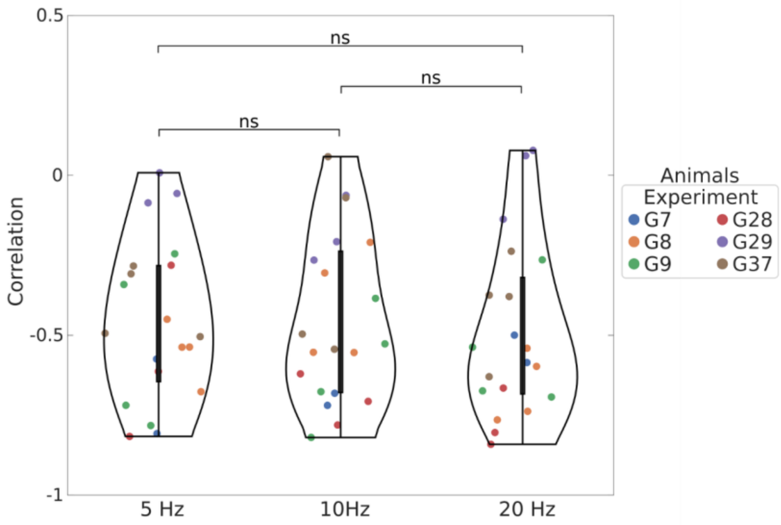


**Supplementary Fig. 8: Effect of different stimulation strengths on correlation between two GABA clusters.** For each experiment and for each stimulation at 5, 10, and 20 Hz, we identified two clusters, calculated the mean activity of each cluster separately, and then computed the correlation between the two clusters within the same experiment. Applying a non-parametric Kruskal-Wallis statistical test shows that there are no significant differences for 5, 10 and 20 Hz stimulation on the correlation of the two clusters. (ns: non-significant).

**Modeling MePD circuit**

To model the interactions between glutamate and GABA neuronal populations in the MePD we are building on our previously proposed network model of the circuit[3]. The model is based on Wilson-Cowan framework and in our case describes the dynamic evolution of calcium activity in neuronal populations due to functional interactions within a synaptically coupled neuronal network.

The model consists of three ordinary differential equations depicting the averaged activity in the neuronal populations of interest, and is governed by the following functions:

$$\frac{G_l}{dt} = \delta(-G_l + (1 - G_l)\varphi(a_l,\, F_l(G_l,\, G_i,\, G_e), \theta_l))$$

$$\frac{dG_i}{dt} = \delta(-G_i + (1 - G_i)\varphi(a_i,\, F_i(G_l,\, G_i,\, G_e), \theta_i))$$

$$\frac{dG_e}{dt} = \delta(-G_e + (1 - G_e)\varphi(a_e,\, F_e(G_l,\, G_i,\, G_e), \theta_e))$$

where $G_l, G_i, G_e$ represent the activity in the populations of glutamatergic neurons (glut), GABA interneurons (GABA$^{int}$) and GABA efferent (GABA$^{eff}$) neurons at time $t$, respectively. The parameter $\delta$ is the time-scaling factor. Function $\varphi$ is a sigmoid stimulus–response function which controls the mean level of activity and given by the formula:

$$\varphi(a,\, F, \theta) = \frac{1}{1 + exp\,(-a(F - \theta))} - \frac{1}{1 + exp\,(a\theta)}$$

where $a$ and $\theta$ define the value of the maximum slope and half-maximum firing threshold, respectively. In the equation, $F$ is the input to the corresponding population, given by the linear sum of excitatory and inhibitory contributions, as follows:

$$F_l(G_l,\, G_i,\, G_e) = (1 - \beta_2)G_l c_{ll} - (1 - \beta_1)G_i c_{il} - (1 - \beta_1)G_e c_{el} + U(t)$$

$$F_i(G_l,\, G_i,\, G_e) = (1 - \beta_2)G_l c_{li} - (1 - \beta_1)G_i c_{ii} - (1 - \beta_1)G_e c_{ei} + U(t) + S(t) + b_i$$

$$F_e(G_l,\, G_i,\, G_e) = (1 - \beta_2)G_l c_{le} - (1 - \beta_1)G_i c_{ie} - (1 - \beta_1)G_e c_{ee} + S(t)$$

where the parameters $c$ represent the strength of interaction from one population to another. The first letter of subscript identifies the population the interaction is coming from, and the second subscript signifies the population the input is going to. The terms $(1 - \beta_1)$ and $(1 - \beta_2)$ represent the suppression of the interaction in populations of GABA neurons and glutamatergic neurons, respectively, with $\beta_1$ and $\beta_2$ as parameters of the proportion of the suppressed interaction in GABA and glutamate neuronal populations, respectively. The term $U$ represents dynamic input from UCN3 neurons to the populations of GABA interneurons and glutamatergic neurons and $S$ is the term responsible for the input to GABA populations during stimulation. The parameter $b_i$ is the baseline activity in GABA interneuron population.

We introduce input from UCN3 neurons into the populations of glutamatergic neurons and GABA interneurons with a strength defined via the parameter $\gamma$. The parameter $\gamma = \gamma_l$ weighs the strength of input from UCN3 neurons to glutamatergic neuronal population and $\gamma = \gamma_e$ for the UCN3 input to the population of GABA interneurons, respectively. The UCN3 input is approximated by a sinusoidal function:

$$U(t) = U_{stim}(t) = \gamma B + Asin\,(2\pi f t)\ ,$$

where $B$ and $A$ correspond to the strength of tonic and periodic input, respectively, while the parameter $f$ is the frequency of the sinusoidal wave. We approximate input from UCN3 neurons during restraint stress using two exponential functions of the form:

$$U(t) = U_{stress}(t) = \gamma(at\,exp\,(-r_2 t) + b(1 - exp\,(-r_1 t)\,)\,),$$

where the function initially increases with the magnitude $a$ then decreases to the plateau valued at $b$. The parameters $r_1$ and $r_2$ are responsible for the rate of increase of the first term and rate of decrease to plateau in the second term.

We mimic the effects of optogenetic stimulation of MePD GABA neuronal populations by introducing periodic input to the variables $G_i$ and $G_e$ as following:

$$S(t) = C + Dsin\,(2\pi f t)\ ,$$

where $C$ and $D$ are baseline excitatory input and the oscillations amplitude, respectively. The parameter $f$ is the frequency of oscillations. The model parameters are given in Supplementary Table 1.

**Supplementary Table 3: MePD model parameter values.**

| Parameter | Description | Value | Reference |
|---|---|---|---|
| $c_{ll}$ | Glutamatergic self-excitation strength [a.u.] | 16 | Derived |
| $c_{li}$ | Interaction strength glut to GABA$^{int}$ [a.u.] | 0 | Derived |
| $c_{le}$ | Interaction strength glut to GABA$^{eff}$ [a.u.] | 11 | Derived |
| $c_{il}$ | Interaction strength GABA$^{int}$ to glut [a.u.] | 0 | Derived |
| $c_{ii}$ | GABA interneurons self-inhibition strength [a.u.] | 30 | Derived |
| $c_{ie}$ | Interaction strength GABA$^{int}$ to GABA$^{eff}$ [a.u.] | 17 | Derived |
| $c_{el}$ | Interaction strength GABA$^{eff}$ to glut [a.u.] | 16 | Derived |
| $c_{ei}$ | Interaction strength GABA$^{eff}$ to GABA$^{int}$ [a.u.] | 15 | Derived |
| $c_{ee}$ | GABA efferents self-inhibition strength [a.u.] | 0 | Derived |
| $\delta$ | Temporal scaling factor [min$^{-1}$] | 3 | Derived |
| $a_l$ | Maximum slope of glut [a.u.] | 1.3 | Derived |
| $a_i$ | Maximum slope of GABA$^{int}$ [a.u.] | 2 | [3] |
| $a_e$ | Maximum slope of GABA$^{eff}$ [a.u.] | 2 | [3] |
| $\theta_l$ | Half-maximum firing threshold for glut [a.u.] | 4 | [3] |
| $\theta_i$ | Half-maximum firing threshold for GABA$^{int}$ [a.u.] | 3.7 | [3] |
| $\theta_e$ | Half-maximum firing threshold for GABA$^{eff}$ [a.u.] | 3.7 | [3] |
| $\beta_1$ | GABAergic interaction suppression coefficient [dimensionless] | 0.5 | [3] |
| $\beta_2$ | Glutamatergic interaction suppression coefficient [dimensionless] | 0.5 | [3] |
| $B$ | Baseline UCN3 neuron activity during stimulation [a.u.] | 2.6 | Derived |
| $A$ | Input from UCN3 neurons [a.u.] | 0.5 | Derived |
| $f$ | Frequency of the sinusoidal wave [sec$^{-1}$] | 10 | Derived |
| $C$ | Baseline excitatory input to GABA population during its stimulation [a.u.] | 8.15 | Derived |
| $D$ | Amplitude of oscillations of GABA stimulation input [a.u.] | 4 | Derived |
| $a$ | Initial growth magnitude during stress [a.u.] | 10 | Derived |
| $b$ | Plateau during restraint stress [a.u.] | 3.9 | Derived |
| $r_1$ | Increase rate parameter during restraint stress [a.u.] | 10 | Derived |
| $r_2$ | Decrease rate parameter during restraint stress [a.u.] | 1 | Derived |
| $\gamma_l$ | Strength of UCN3 input to glut [a.u.] | 1 | Derived |
| $\gamma_i$ | Strength of UCN3 input to GABA$^{int}$ [a.u.] | 0.115 | Derived |
| $b_i$ | GABA interneuron baseline activity [a.u.] | 9 | Derived |

**Coarse-grained model of ARC KNDy network**

We use our previously established model of ARC KNDy network[4] to investigate the effects of MePD interventions on the GnRH pulse generator. The model is given by the system of three ordinary differential equations:

$$\frac{dD}{dt} = f_D(v) - d_D D,$$

$$\frac{dN}{dt} = f_N(N, v) - d_N N,$$

$$\frac{dv}{dt} = f_v(N, v) - d_v v,$$

where $D$ and $N$ represent the concentration of dynorphin and neurokinin B produced by the population, $v$ and describes the averaged firing activity in the population in spikes per minute. The terms $d_D, d_N$ and $d_v$ stand for the linear decay for each variable. The secretion rates for dynorphin and neurokinin are given by the function $f_D$ and $f_N$:

$$f_D(v) = k_D \frac{v^2}{v^2 + K_{v,1}^2},$$

$$f_N(N, v) = k_N \frac{v^2}{v^2 + K_{v,2}^2} \frac{K_D^2}{K_D^2 + D^2},$$

where $k_D$ and $k_N$ signify the neuropeptides' secretion rates; $K_{v,1}$and $K_{v,2}$ describe the frequency value for which the rate of dynorphin and neurokinin B secretion is half-maximum; and $K_D$ describes the dynorphin concentration that results in half-maximum inhibition. The original function $f_v$ has been modified as per Nechyporenko *et al*[3]:

$$f_v = v_0 \frac{1}{1 + exp\ (k(-I + m))},$$

where $m$ defines half-maximum level of synaptic input and $k$ is the membrane's time constant, which determines how quickly the neuron's membrane potential changes in response to inputs. The parameter $v_0$ is the maximum increase to synaptic inputs $I$ [Hz], which is defined as following:

$$I = I_0 + p_v \frac{N^2}{N^2 + K_N^2} v - jG_e,$$

where $I_0$ is the basal input to the population, $p_v$ and $K_N$ are neurokinin B's half-maximal effect and the positive-feedback strength, respectively. The term $jG_e$ signifies MePD GABAergic input into KNDy with the pre-synaptic firing rate conversion parameter. The KNDy network model parameters are given in Supplementary Table 2.

**Supplementary Table 4: KNDy model parameter values.**

| Parameter | Description | Value | Reference |
|---|---|---|---|
| $d_D$ | Dynorphin degradation rate [$min^{-1}$] | 0.2 | [4] |
| $d_N$ | Neurokinin B degradation rate [$min^{-1}$] | 1 | [4] |
| $d_v$ | Firing rate reset rate [$min^{-1}$] | 10 | [5] |
| $k_D$ | Dynorphin signaling strength [nM $min^{-1}$] | 4 | [6] |
| $k_N$ | Neurokinin B signaling strength [nM $min^{-1}$] | 40 | [6] |
| $p_v$ | Effective strength of synaptic input [a.u.] | 0.008 | [6] |
| $v_0$ | Maximum rate of neuronal activity increase [spikes $min^{-2}$] | 25 000 | [7] |
| $K_D$ | Dynorphin $IC_{50}$ [nM] | 0.3 | [7] |
| $K_N$ | Neurokinin B $IC_{50}$ [nM] | 4 | [8] |
| $K_{v,1}$ | Firing rate for half-maximal dynorphin secretion [spikes $min^{-1}$] | 600 | [9] |
| $K_{v,2}$ | Firing rate for half-maximal neurokinin B secretion [spikes $min^{-1}$] | 200 | [9] |
| $k$ | Membrane's time constant [min] | 10 | [3] |
| $m$ | Half-maximal firing rate [$min^{-1}$] | 0.5 | [3] |
| $I_0$ | Basal activity [Hz] | 0.2 | Fixed |
| $j$ | pre-synaptic firing rate conversion parameter for GABAergic projections [Hz] | 0.5 | Fixed |

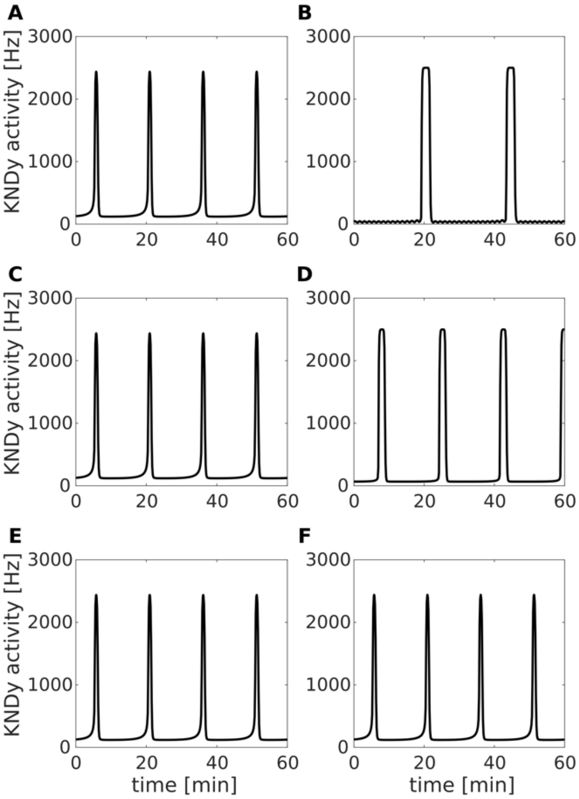


**Supplementary Fig. 9: Model calibration.** (**A**) KNDy activity with no interventions (control) has an inter-pulse interval (IPI) of 15.18 min. (**B**) During simulation of the UCN3 stimulation IPI increases to 24.22 min. (**C**) Simulation of the effects of GABA receptor antagonist (suppression of GABAergic interactions) does not affect the KNDy IPI (15.18 min) (**D**) Simulation of the combined effects of stimulation of UCN3 and GABA receptor antagonist changes KNDy IPI to 17.32 min. (**E**) Simulation of the effects of glutamate receptor antagonist and (**F**) glutamate receptor antagonist together with stimulation of UCN3 does not affect KNDy IPI (15.18 min and 15.31 min, respectively).

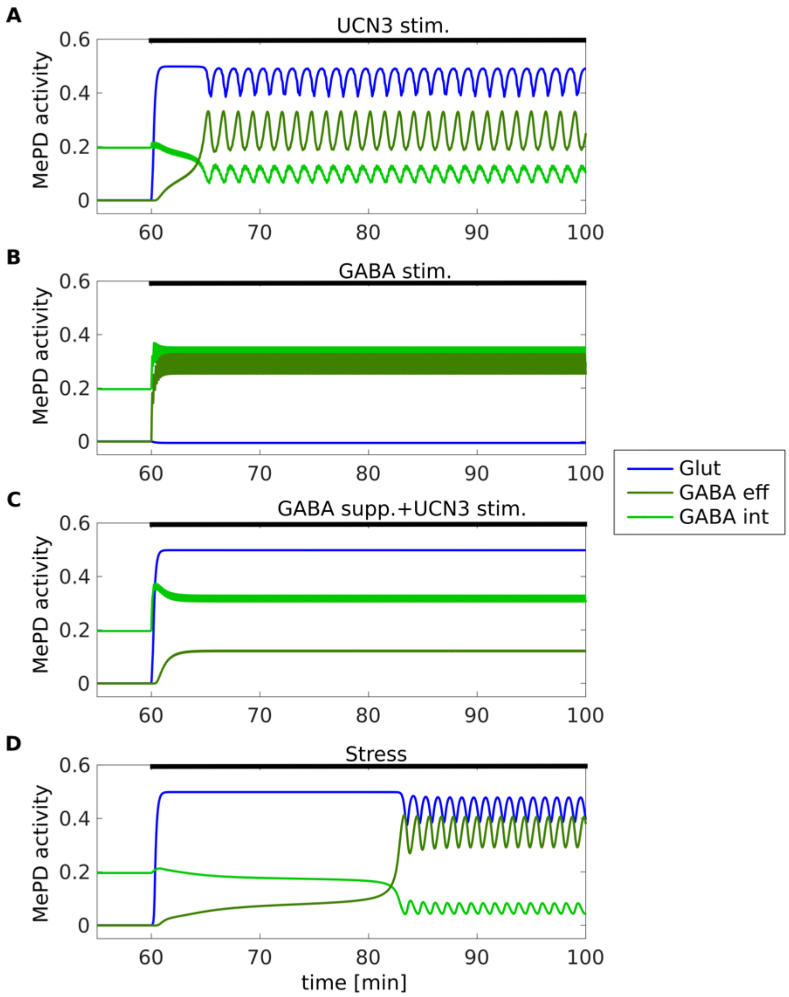


**Supplementary Fig. 10: MePD populations activity during interventions.** Activity of glutamate neurons, GABA interneurons and GABA efferent neurons during (**A**) stimulation of UCN3 neurons, (**B**) stimulations of GABA neurons, (**C**) stimulation of UCN3 neurons and suppression of GABA neurons, (**D**) restraint stress.

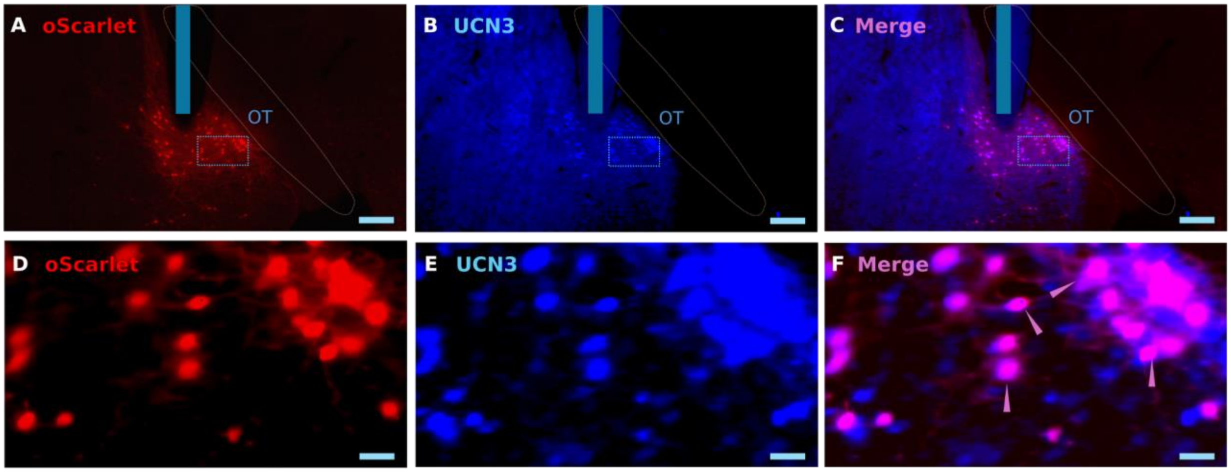


**Supplementary Figure 11**：**Validation of AAV-nEF-Con/Foff-ChRmine-oScarlet expression in UCN3 neurons in the MePD.** (**A**&**D**) Red fluorescent labeled UCN3 neurons expressing ChRmine-oScarlet. (**B**&**E**) Blue fluorescent labeled UCN3 neurons immuno-stained positive for UCN3 antibody tagged with Alexa Fluor 405. (**C**&**F**) The merged images (magenta) demonstrate the co-localization of Con/Foff ChRmine-oScarlet and UCN3 immunoreactivity. The blue vertical bars in **A** to **C** indicate the position of the fibre optic cannula. The pink arrowheads indicate double-labeled neurons. Scale bars represent (**A** to **C**) 200 μm, (**D** to **F**) 25 μm. OT, optic track.

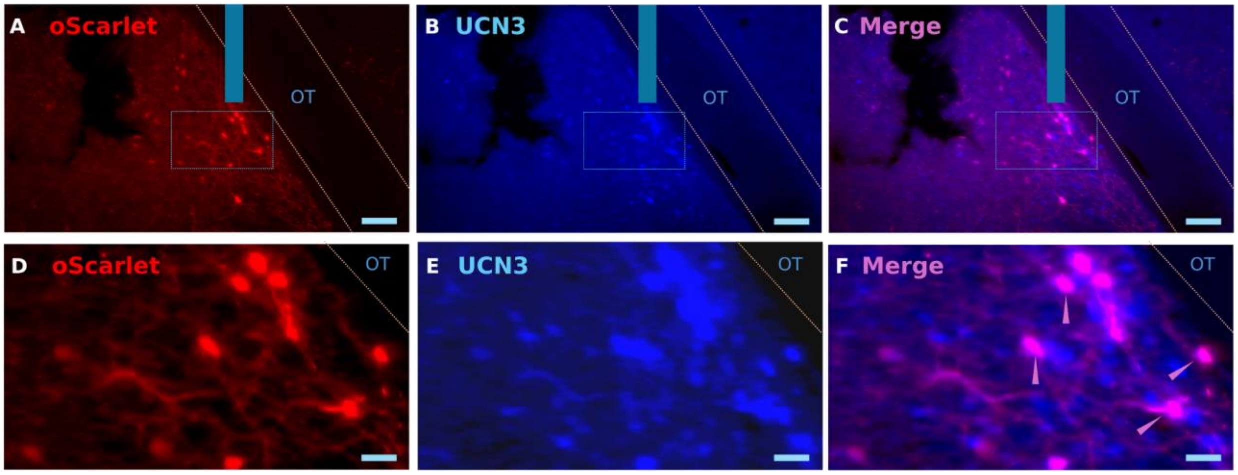


**Supplementary Figure 12：Validation of AAV-nEF-Con/Fon-ChRmine-oScarlet expression in UCN3 neurons in the MePD.** (**A**&**D**) Red fluorescent labeled UCN3 neurons expressing ChRmine-oScarlet. (**B**&**E**) Blue fluorescent labeled UCN3 neurons immunopositive for the UCN3 antibody tagged with Alexa Fluor 405. (**C**&**F**) The merged images (magenta) demonstrate the co-localization of Con/Fon ChRmine-oScarlet and UCN3 immunoreactivity. The blue vertical bars in **A** to **C** indicate the position of the fibre optic cannula. Pink arrowheads indicate double-labeled neurons. Scale bars represent (**A** to **C**) 100 μm, (**D** to **F**) 25 μm. OT, optic track.

**Supplementary Table 5: LH pulse amplitude and meal levels before and during simultaneous optogenetic stimulation in UCN3-Cre::VGAT-Flpo female mice.** The LH pulse amplitude and mean LH levels of the UCN3-Cre::VGAT-Flpo female mice injected with intersectional virus in the MePD did not significantly change after optic stimulation or control optic stimulation (two-way ANOVA). Data are presented as mean ± SEM.

| Group | Mean LH level (ng/mL) | | LH pulse amplitude (ng/mL) | |
|---|---|---|---|---|
| | Before | After | Before | After |
| Con/Foff virus + stimulation | 4.08 ± 0.33 | 4.20 ± 0.54 | 2.83 ± 0.72 | 3.61 ± 0.72 |
| Con/Foff without stimulation | 4.02 ± 0.35 | 3.95 ± 0.18 | 3.42 ± 0.36 | 3.00 ± 0.40 |
| Con/Foff control virus + stimulation | 4.79 ± 0.60 | 4.67 ± 0.66 | 3.09 ± 0.35 | 3.24 ± 0.59 |
| Con/Fon virus + stimulation | 4.80 ± 0.26 | 5.00 ± 0.26 | 3.75 ± 0.49 | 3.92 ± 0.62 |
| Con/Fon without stimulation | 5.35 ± 0.78 | 5.62 ± 0.84 | 4.34 ± 0.61 | 4.51 ± 0.31 |
| Con/Foff+Coff/Fon virus + stimulation | 4.68 ± 0.25 | 4.80 ± 0.43 | 2.88 ± 0.39 | 3.51 ± 0.51 |
| Con/Foff+Coff/Fon without stimulation | 3.74 ± 0.34 | 4.22 ± 0.63 | 3.79 ± 0.48 | 3.67 ± 0.73 |

**Supplementary Table 6: LH pulse amplitude and mean levels before and during simultaneous optogenetic stimulation in VGAT-Cre-tdTomato female mice.** The mean LH levels of the VGAT-Cre-tdTomato female mice injected with cre-dependent virus expressing ChR2 is lower in the post-stimulation period compared with pre-stimulation period and the control groups. The symbols # and † indicate p <0.01 and p <0.05, respectively, vs the group of GABA neuron virus + stimulation (post-stimulation period) (two-way ANOVA, Tukey’s post-hoc). Data are presented as mean ± SEM.

| Group | Mean LH level (ng/mL) | | LH pulse amplitude (ng/mL) | |
|---|---|---|---|---|
| | Before | After | Before | After |
| GABA neuron virus + stimulation | 2.35 ± 0.15 † | 1.22 ± 0.22 | 3.38 ± 0.41 | 2.83 ± 0.21 |
| GABA neuron virus without stimulation | 2.27 ± 0.28 † | 2.42 ± 0.12 † | 3.48 ± 0.48 | 3.41 ± 0.41 |
| GABA neuron control virus + stimulation | 2.96 ± 0.28 # | 2.99 ± 0.30 # | 2.07 ± 0.32 | 2.42 ± 0.34 |
